# Delicate tuning of H_2_O_2_ by salicylic acid, catalase 2, and AtSAP5 incites plant immunity

**DOI:** 10.1101/2024.09.30.615974

**Authors:** Li Chang, Yi-Shu Chiu, Shang-Huan Chiang, An-Po Cheng, Yuh Tzean, Hsin-Hung Yeh

## Abstract

Salicylic acid (SA) triggers plant immunity through redox state changes, initially oxidizing and then reducing. SA binds and inhibits the H_2_O_2_-scavenging enzyme catalase 2 (CAT2), promoting an initial oxidative state. However, a reduced state is crucial for a robust immune response. In *Arabidopsis*, the SA-induced A20/AN1 zinc finger protein, AtSAP5, competes with SA for CAT2 binding, maintaining CAT2 activity. This reduces H_2_O_2_ levels and prevents CAT2 degradation through inhibition of H_2_O_2_-induced autophagy formation, which forms a positive feedback loop to CAT2 accumulation. Consequently, cells shift to a more reduced state, activating immune responses. Our findings reveal a nuanced regulatory mechanism through which H_2_O_2_ levels are controlled by SA with cross-kingdom conserved proteins AtSAP5 and CAT2, managing redox states and plant immunity.

## INTRODUCTION

Salicylic acid (SA) is an important phytohormone that regulates plant immunity to combat biotrophic and hemi-biotrophic pathogen infection (*1, 2*). The *NONEXPRESSER OF PATHOGENESIS-RELATED GENES 1* (*NPR1*) has been identified as the key regulatory gene in SA-mediated immune response (*3-5*). NPR1 serves as an SA binding receptor and functions as a transcriptional mediator to regulate the expression of about 90% of SA-mediated immune response genes (*1, 2, 6-8*). However, the transcription level of *NPR1* is only induced 2–3 fold upon pathogen infection or SA treatment (*9, 10*), and the post-translational modification (PTM) of NPR1 has been demonstrated to play an important role in the regulation of SA-responsive immune genes (*11-13*). Several PTMs of NPR1 have been reported, including the reduction of the NPR1 oligomer into monomers, phosphorylation, and sumoylation (*11*). The dynamic PTM of NPR1 enables dedicated control of immune induction with minimal adverse effects on plant growth (*14*). In a healthy plant, NPR1 exists in the cytosol as an oligomer generated through the formation of intermolecular disulfide bonds (*12*). Pathogen infection or exogenous application of SA induces the synthesis of SA (*15*). SA induces a reduced state that facilitates NPR1 monomerization, which is critical for NPR1 nucleus translocation and subsequently for the expression of its own and the activation of massive downstream SA-responsive immune genes (*12, 16*).

Indeed, pathogen infection or INA treatment (an SA analog) induces a biphasic change in the redox state in *Arabidopsis*, first oxidative and then reducing, as determined by measuring the ratio of reduced/oxidized glutathione (*12*). The initial oxidate state was suggested to serve as a signal for triggering plant immunity (*17*). It is known that SA binds to H_2_O_2_-scavenging enzymes, catalase (CAT), and ascorbate peroxidase (APX), in tobacco, and that leads to the accumulation of H_2_O_2_, SA free radicals, and other reactive oxygen species (ROS) (*18-23*). In *Arabidopsis*, SA also inhibits CAT2. The CAT isoform contributes 90% of the catalase activity in leaves (*24*), which results in increased H_2_O_2_ accumulation (*25*). This may contribute to the initial SA-induced oxidative state in *Arabidopsis* (*12*). However, a more reduced state is necessary for the nuclear translocation of NPR1 to activate plant immunity. These findings raise the important but unsolved question of how plants transition from the SA-induced oxidative state to a more reduced state to activate plant immunity.

Proteins containing A20 and/or AN1 zinc finger domains are conserved across kingdoms (*26*). In plants, these A20 and/or AN1 proteins are named stress-associated proteins (SAPs), which were initially characterized as being involved in tolerance against abiotic stresses (*27*). However, more and more studies have indicated that SAPs also play an important role in immunity against biotic stresses. SAPs from rice (OsSAP1), tomato (SISAP3 and SISAP4), *Arabidopsis* (AtSAP5, AtSAP9, and AtSAP12), and *Phalaenopsis* orchid (Pha13 and Pha21) have been reported to participate in immunity in response to different pathogens including bacteria, fungi, nematodes, and viruses (*28-34*).

Our previous study indicated that SAPs, including *Phalaenopsis* orchid Pha13 and Pha21 and *Arabidopsis* AtSAP5, are important hubs in SA-mediated plant immunity (*32*). Pha13 and Pha21 coordinate the activation of antiviral immune genes, including SA-regulated *NPR1-* dependent, *NPR1*-independent, and RNAi-related genes (*32, 34*). The *Arabidopsis* homolog of Pha13 and Pha21, AtSAP5, has a similar role in virus resistance as well as activation of SA-regulated and RNAi-related antiviral immune genes (*32, 34, 35*). In addition, the tomato SAPs (SlSAPs) also contribute to plant immunity through different plant hormone-mediated immune pathways. SlSAP3 is involved in SA-mediated immune response gene expression and enhances protection against the biotrophic pathogen *Pseudomonas syringae* pv. *tomato* DC3000 (*Pst* DC3000) (*31*). SlSAP4 functions in the jasmonate-and ethylene-mediated pathways to enhance immunity against the necrotrophic pathogen *Botrytis cinerea* (*29*). SlSAP12 and its *Arabidopsis* homolog AtSAP12 have been reported to be targeted by an effector, disulphide isomerase-like protein, encoded by *Meloidogyne incognita*, which is necessary for disease establishment in both tomato and *Arabidopsis* (*33*). Overexpression of rice OsSAP1 in tobacco enhances plant resistance to bacterial pathogens (*36*). This indicates that SAP-mediated plant immunity is conserved among plants including both dicots and monocots. Although the importance of SAPs in plant immunity has been demonstrated, the underlying mechanism remains unknown.

In this study, we demonstrate that the SA-induced AtSAP5 functions as a redox modulator of CAT2 in H_2_O_2_-scavenging activity. AtSAP5 binds to CAT2 and counteracts the negative effect of SA on CAT2 H_2_O_2_-scavenging activity. The decreased H_2_O_2_ prevents the induction of autophagy, thus preventing the degradation of CAT2 through the autophagy pathway, which forms a positive feedback loop to increase the CAT2 accumulation and decrease H_2_O_2_. This promotes cells to gradually shift to a more reduced state, which is important for NPR1 nuclear translocation and activation of downstream immune response genes. These data shed new light on the mechanism through which SA, CAT2, and AtSAP5 regulate the redox state to activate plant immunity.

## Results

### AtSAP5-mediated plant immunity against *Pst* DC3000 is mainly dependent on NPR1 in

#### Arabidopsis

To analyze whether NPR1 is involved in AtSAP5-mediated plant immunity, we inoculated the following lines with *Pst* DC3000: wild-type (WT) *Arabidopsis* (Col-0), transgenic *Arabidopsis* overexpressing AtSAP5 (AtSAP5-oe-11 and AtSAP5-oe-12), the *npr1* mutant, and two lines of transgenic *Arabidopsis* overexpressing AtSAP5 in the *npr1* mutant background (AtSAP5-oe/*npr1*-1 and AtSAP5-oe/*npr1*-2) (Fig. 1A).

**Fig. 1.**
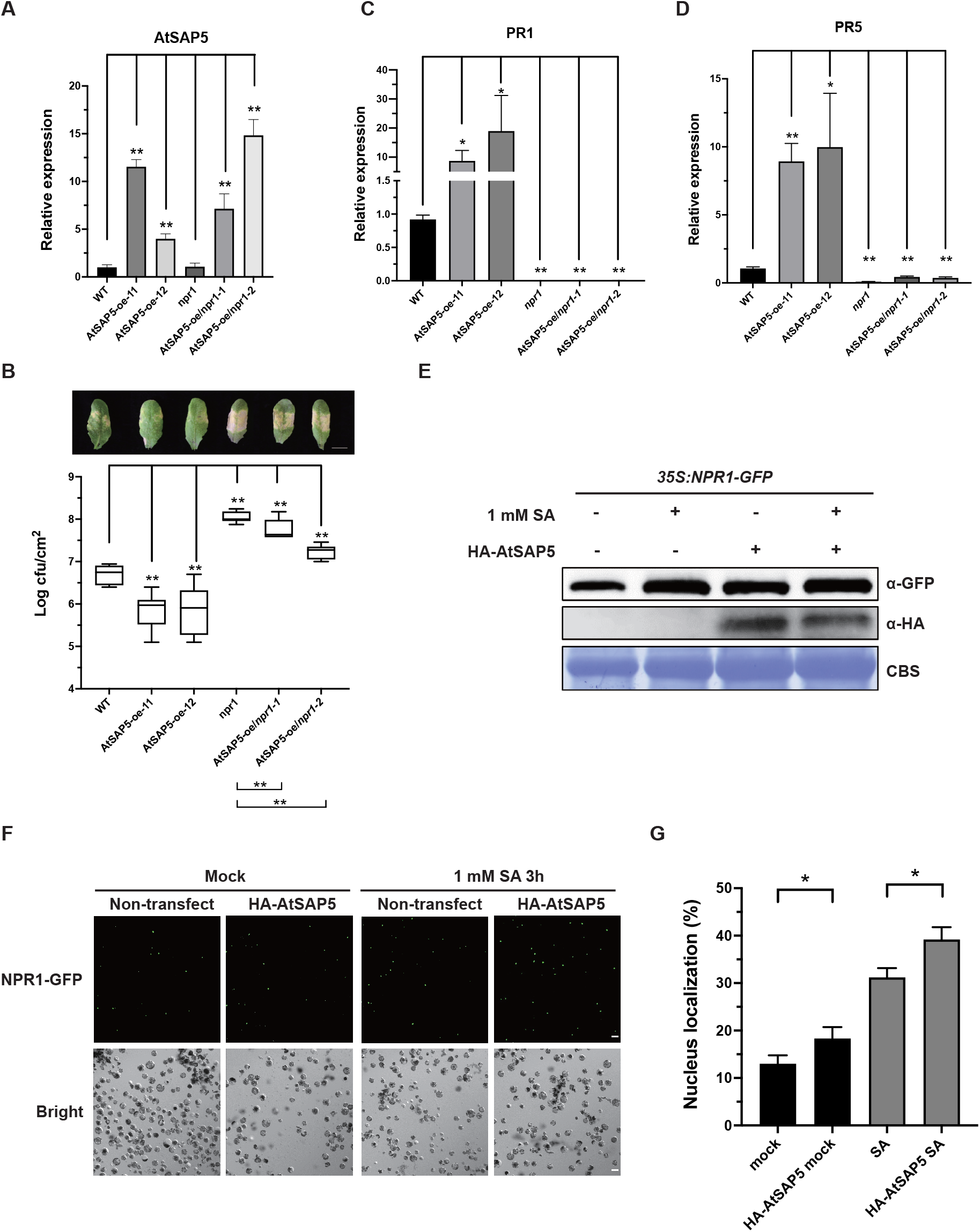
NPR1 is involved in AtSAP5-mediated plant immunity against *Pseudomonas syringae* pv. tomato DC3000 (*Pst* DC3000) and AtSAP5 promotes NPR1 nuclear translocation in *Arabidopsis*. Wild-type (WT) *Arabidopsis* (Col-0), transgenic *Arabidopsis* overexpressing AtSAP5 (AtSAP5-oe-11 and AtSAP5-oe-12), *NPR1* mutant (*npr1*), and transgenic *Arabidopsis* overexpressing AtSAP5 in *npr1* (AtSAP5-oe/*npr1*-1 and AtSAP5-oe/*npr1*-2) were used in the analysis. (**A**) The RNA level of AtSAP5 of each plant was analyzed by RT-qPCR. Data represent mean ± SD; n = 4 to 7 biological replicates. (**B**) The bacterial accumulation level and symptoms on leaves of each plant inoculated with *Pst* DC3000 at 3 days post-inoculation were analyzed. Scale bar, 1 cm. n = 8 biological replicates. (**C** and **D**) The relative expression levels of immune marker genes *PR1* and *PR5* in flagellin 22-treated plants were analyzed by RT-qPCR at 24 h post-treatment. Data represent mean ± SD; n = 5 biological replicates. (**A** to **D**) *, *P* < 0.05, **, *P* < 0.01, Student’s t-test. The *AtActin* was used as an internal control for RT-qPCR normalization. Three independent experiments showed similar results. **(E** to **G)** The effect of AtSAP5 on NPR1 nucleus localization was analyzed. Protoplasts were isolated from a transgenic *Arabidopsis* line overexpressing *NPR1-*green fluorescent protein (GFP) in the *npr1* background (NPR1-GFP/*npr1*). The protoplasts were transfected with vectors expressing AtSAP5-fused with HA (HA-AtSAP5). After 16 h of incubation, the cells were treated with H_2_O (Mock) or SA (1 mM). **(E)** The expression of HA-AtSAP5 and NPR1-GFP were detected by immunoblotting using anti-HA and anti-GFP antibodies, respectively. Coomassie brilliant blue staining (CBS) was used as a loading control. **(F)** The indicated fluorescence was detected by confocal microscopy. Representative confocal microscopy images are shown. The GFP signal indicates the NPR1-GFP protein. **(G)** The ratio of NPR1-GFP signal translocated into the nucleus was analyzed by dividing the number of GFP signals by the total number of cells in a single field of view. Data represent mean ± SD. The results were combined from 3 independent experiments, and each experiment result was the average of three different confocal microscopy fields of view; *, *P* < 0.05, Student’s t-test.

Both AtSAP5-oe-11 and AtSAP5-oe-12 showed enhanced resistance to *Pst* DC3000 as compared with WT (Col-0) (Fig. 1B); however, *npr1*, AtSAP5-oe/*npr1*-1, and AtSAP5-oe/*npr1*-2 all showed compromised immunity in comparison to WT (Col-0), AtSAP5-oe-11 and AtSAP5-oe-12 (Fig. 1B).

In addition, we also analyzed the expression level of two known NPR1-dependent marker genes, *PR1* and *PR5* (*37, 38*), in the abovementioned plants at 24 h post-treatment with flagellin 22 (flg22), a known pathogen-associated molecular pattern that can trigger plant immunity (*39, 40*). Enhanced expression of *PR1* and *PR5* was observed in both AtSAP5-oe-11 and AtSAP5-oe-12 as compared with WT (Col-0) (Fig. 1C and 1D); however, reduced expression of *PR1* and *PR5* was observed in *npr1*, AtSAP5-oe/*npr1*-1, and AtSAP5-oe/*npr1*-2 (Fig. 1C and 1D).

Collectively, our data indicate that AtSAP5-mediated plant immunity largely depends on NPR1.

### Overexpression of AtSAP5 enhances the nuclear translocation of NPR1

NPR1 nucleus translocation is important for the activation of SA-mediated immune genes (*1, 12*). Therefore, we analyzed whether AtSAP5 affects NPR1 nucleus translocation. We generated *Arabidopsis* lines overexpressing NPR1 fused with green fluorescent protein (GFP) in the *npr1-1* mutant background (*35S::NPR1-GFP*/*npr1*) as reported previously (*12*). Protoplasts isolated from *35S::NPR1-GFP*/*npr1* were used for transfection assays. AtSAP5 fused with HA (HA-AtSAP5) was transfected into *35S::NPR1-GFP*/*npr1* protoplasts under mock (H_2_O) or 1 mM SA treatment. The expression of HA-AtSAP5 in cells was confirmed by immunoblotting using an anti-HA antibody (Fig. 1E).

In the non-transfected cells, SA treatment increased the ratio of cells showing the nuclear NPR1-GFP signal (31.2%) as compared with mock-treated cells (13%) (Fig. 1F and 1G), which is similar to the findings of a previous report (*12*). In addition, more NPR1-GFP signals were observed in cells overexpressing HA-AtSAP5 (18.3%) than in the non-transfected cells (13%) under mock treatment (Fig. 1F and 1G). In addition, more NPR1-GFP signals were also observed in cells overexpressing HA-AtSAP5 (39.2%) than in the non-transfected control (31.2%) under SA treatment (Fig. 1F and 1G). The results indicate that AtSAP5 facilitates NPR1 nucleus translocation.

### AtSAP5 interacts with CAT2

To investigate how AtSAP5 regulates SA-mediated immune response genes, we performed an immunoprecipitation (IP) assay to identify putative AtSAP5-interacting proteins in *Arabidopsis*. Total proteins were extracted from leaves of WT (Col-0) and transgenic *Arabidopsis* lines overexpressing AtSAP5 (AtSAP5-oe-11 and AtSAP5-oe-12), and anti-AtSAP5 antibody was used to pull down the AtSAP5-interacting proteins. The protein complexes pulled down by IP were further analyzed by liquid chromatography-tandem mass spectrometry (LC-MS-MS) (Table S1).

CAT2 was among the putative interactors showing 32, 42, and 39 peptide-spectrum match (PSM) in WT (Col-0), AtSAP5-oe-11 and AtSAP5-oe-12 plants, respectively (Table S1). CAT2 serves as the major CAT isoform in the leaves of *Arabidopsis* (*24*) and was reported to be involved in SA-mediated responses (*1, 25, 41-43*). Therefore, we selected CAT2 for further analysis. An *in vitro* pull-down assay was conducted to analyze the interaction between AtSAP5 and CAT2. We were unable to purify the full-length of CAT2-His from *Escherichia coli*; thus, we overexpressed and purified CAT2-His and GFP-His recombinant proteins from *Nicotiana benthamiana* (Fig. S1). AtSAP5-His recombinant proteins purified from *E. coli* were then mixed with CAT2-His or GFP-His, pulled down with anti-AtSAP5 antibody, and then detected by anti-His antibody. The results showed that AtSAP5-His interacts with CAT2-His but not GFP-His (Fig. 2A).

**Fig. 2.**
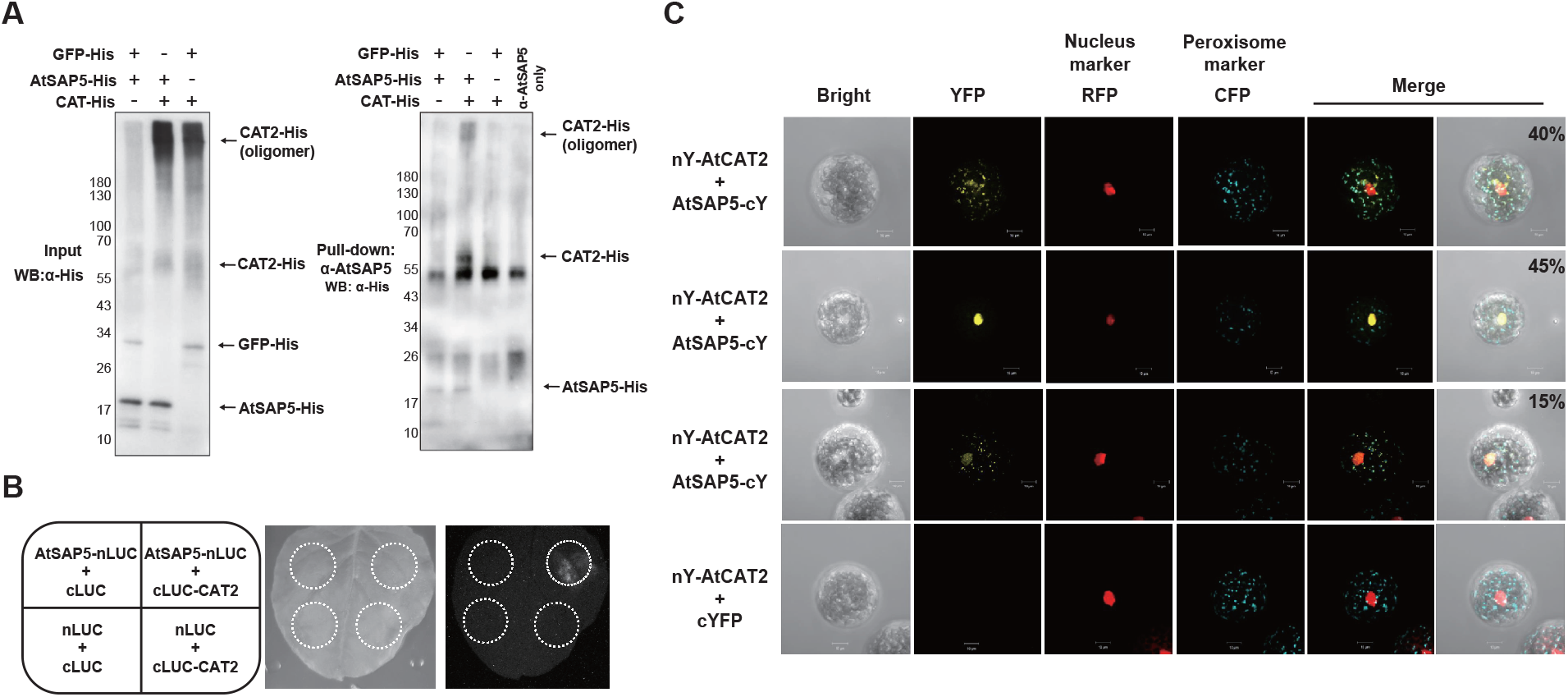
AtSAP5 interacts with CAT2. **(A)** An *in vitro* AtSAP5-pull down assay was performed to detect the interaction between AtSAP5 and CAT2. Two of the three recombinant proteins (GFP-His, AtSAP5-His, and CAT2-His) were mixed and incubated. Pull-down was performed using anti-AtSAP5 antibody. Samples before (input) and after pull-down were analyzed by western blot with anti-His antibody. The anti-AtSAP5 antibody only was used as a pull-down control. GFP-His, CAT2-His, CAT2-His oligomer, and AtSAP5-His are indicated by arrows. **(B)** The split luciferase assay was performed by infiltrating *Agrobacterium* carrying the constructs into *Nicotiana benthamiana* leaves to express the indicated proteins. Three days after infiltration, luciferin was sprayed onto the infiltrated leaves, and chemiluminescence images were taken by a CCD camera (Azure Biosystems). AtSAP5 fused with the N terminus of luciferase (AtSAP5-nLUC) and CAT2 fused with the C terminus of luciferase (cLUC-CAT2) are indicated. nLUC and cLUC were used as controls. **(C)** The bimolecular fluorescence complementation assay was performed by co-expressing AtSAP5 fused with the C terminus of yellow fluorescent protein (cYFP) (AtSAP5-cY) with CAT2 fused with the N terminus of YFP (nYFP) (nY-CAT2) in protoplasts isolated from *Arabidopsis*. cYFP served as a control. The nuclear localization signal-fused red fluorescence protein (RFP) was used as a nucleus marker. The peroxisomal targeting signal 1-fused cyan fluorescent protein (CFP) was used as a peroxisome marker. Representative images were obtained by confocal microscopy. Scale bars represent 10 μm. The ratio indicates the number of cells showing a specific localization pattern among all examined cells with an interaction signal. (**A** to **C**) Three independent experiments showed similar results.

To determine whether the interaction between AtSAP5 and CAT2 occurs in cells, split luciferase and bimolecular fluorescence complementation (BiFC) assays were conducted. In the split luciferase assays, complemented luciferase activity was detected when AtSAP5 fused with the N terminus of luciferase and CAT2 fused with the C terminus of luciferase were co-expressed in leaves of *N. benthamiana* (Fig. 2B). In the BiFC assay, a yellow fluorescent protein (YFP) signal was observed when AtSAP5 fused with the C terminus of YFP (AtSAP5-cY) and CAT2 fused with the N terminus of YFP (nY-CAT2) were co-expressed in protoplasts isolated from WT (Col-0) (Fig. 2C). Of note, the YFP was co-localized with the nucleus and/or peroxisome marker (Fig. 2C). About 45% of cells showed YFP signals in the nucleus only, 40% of cells showed YFP signals in the peroxisome and part of the cytosol, and 15% of cells showed YFP signals in the nucleus,peroxisome, and part of the cytosol (Fig. 2C). Collectively, our data indicates that AtSAP5 interacts with CAT2.

### AtSAP5 does not ubiquitinate CAT2

AtSAP5 has been reported to be an E3 ligase with self-ubiquitination activity (*44, 45*). To analyze whether CAT2 serves as an E3 ligase substrate of AtSAP5, we extracted total proteins from WT (Col-0), AtSAP5-oe-11 and AtSAP5-oe-12, and used anti-CAT antibody to pull down CAT2 protein followed by LC-MS-MS analysis. CAT proteins were the most abundant protein detected from all samples (Table S2). Several ubiquitinated proteins were detected (Table S2), whereas all detected CAT2 showed no ubiquitination signals (Table S2).

In addition, *in vitro* E3 assay was conducted using purified AtSAP5-His and CAT2-His. Self-ubiquitination of AtSAP5 was detected by the use of anti-FLAG antibody in the presence of FLAG-ubiquitin (FLAG-Ub), human E1(hE1) and E2(hE2), and AtSAP5-His (Fig. S2A lane 5). However, no ubiquitinated CAT-His was observed in the presence of FLAG-Ub, hE1 and hE2, AtSAP5-His, and CAT2-His (Fig. S2A lane 6). We also detected CAT2-His using an anti-CAT antibody (Fig. S2B). No obvious molecular weight shift in CAT2-His was observed in the presence of FLAG-Ub, hE1 and hE2, AtSAP5-His, and CAT2-His (Fig. S2B lane 6). These results suggested that CAT2 is not an E3 ligase substrate of AtSAP5.

### AtSAP5 enhances CAT2 protein accumulation and increases H_2_O_2_-scavenging ability *Arabidopsis*

Because AtSAP5 interacts with CAT2, we analyzed the effect of AtSAP5 on H_2_O_2_-scavenging activity within the plant. Increased H_2_O_2_ degradation ability was observed from total proteins extracted from leaves of transgenic *Arabidopsis* overexpressing AtSAP5 (AtSAP5-oe-11, and AtSAP5-oe-12) in comparison to WT (Col-0) (Fig. 3A).

**Fig. 3.**
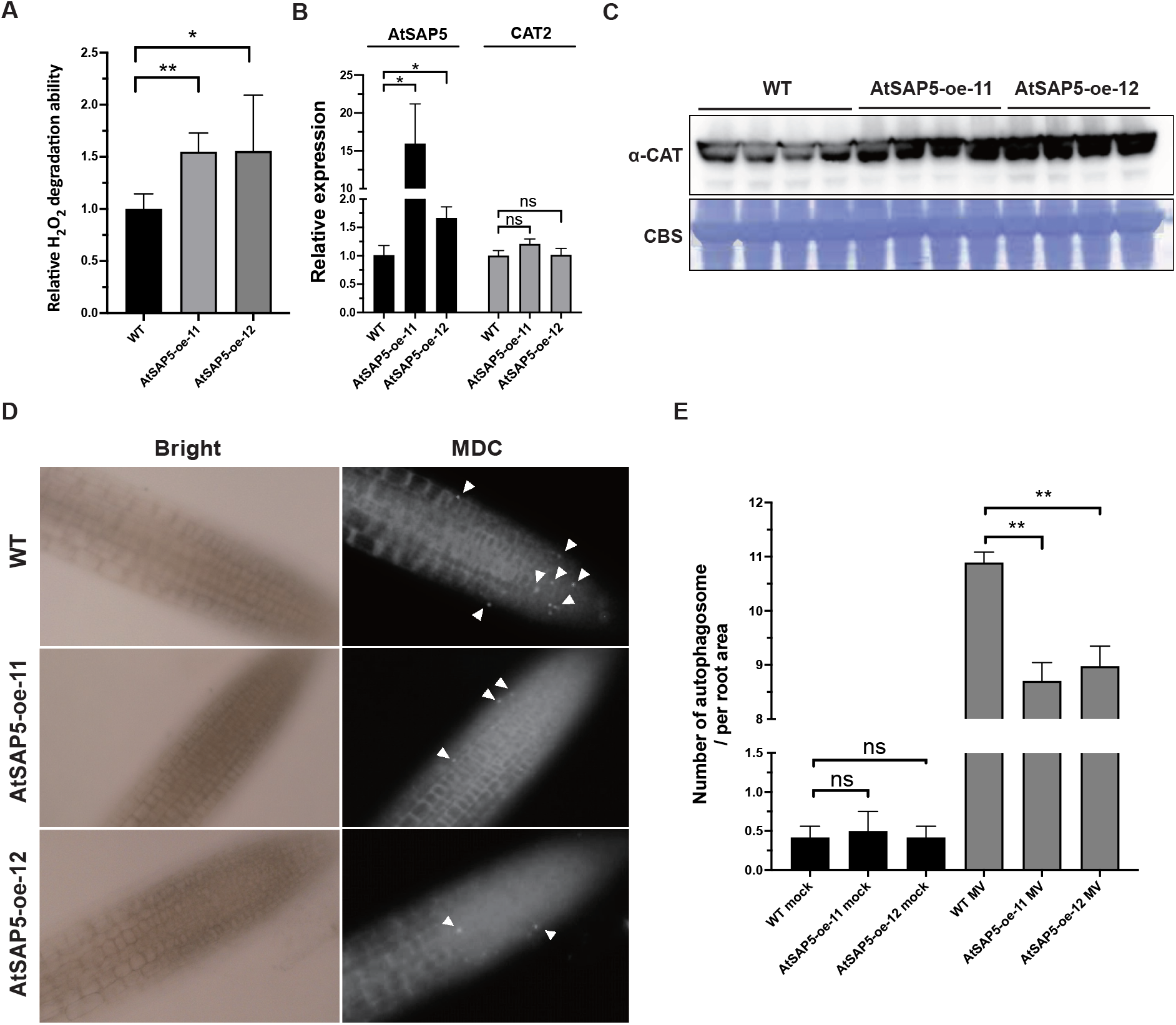
Overexpression of AtSAP5 enhances H_2_O_2_ degradation activity and CAT2 protein accumulation, and reduces the number of autophagosomes in *Arabidopsis*. **(A)** Total protein extracted from leaves of WT (Col-0) and transgenic *Arabidopsis* overexpressing AtSAP5 (AtSAP5-oe-11, and AtSAP5-oe-12) was analyzed for H_2_O_2_ degradation activity. Data represent mean ± SD; n = 5 biological replicates; **, *P* < 0.01; *, *P* < 0.05, Student’s *t*-test. Three independent experiments showed similar results. **(B)** Expression levels of AtSAP5 and CAT2 were analyzed by RT-qPCR from leaves of WT, AtSAP5-oe-11, and AtSAP5-oe-12. Data represent mean ± SD; n = 5 biological replicates; *, *P* < 0.05, Student’s t-test. The *AtActin* was used as an internal control for normalization. Three independent experiments showed similar results. **(C)** The protein level of CAT in WT and transgenic *Arabidopsis* overexpressing AtSAP5 was analyzed by anti-CAT antibody. Coomassie brilliant blue staining (CBS) was used as a loading control. **(D)** The WT, AtSAP5-oe-11, and AtSAP5-oe-12 plants were treated with 10 μm methyl viologen (MV) to induce the formation of autophagosomes. At 2 days post-treatment, the autophagosomes (white arrows) in the roots treated with monodansylcadaverine (MDC) staining were observed using a fluorescence microscope under a DAPI filter. Scale bar, 20 μm. (**E**) The number of autophagosomes in the root area was analyzed in WT, AtSAP5-oe-11, and AtSAP5-oe-12 treated with H_2_O (mock) or 10 μm MV. Data represent mean ± SD. The results from 3 independent experiments were combined; *\*\*, P < 0.01*, Student’s *t* test.

The AtSAP5-mediated enhancement of H_2_O_2_-scavenging ability may result from the increased accumulation of CAT2 protein. Therefore, we analyzed whether AtSAP5 increases the CAT2 protein expression. The CAT2 mRNA accumulation showed no significant difference between the WT (Col-0) and transgenic *Arabidopsis* overexpressing AtSAP5 (Fig. 3B); however, an increased CAT2 protein level was observed in AtSAP5-oe-11 and AtSAP5-oe-12 (Fig. 3C). Because our CAT antibody may not be able to differentiate CAT isoform, we also analyzed the abundance of CAT using anti-CAT antibody to pull down proteins extracted from WT (Col-0),

AtSAP5-oe-11 and AtSAP5-oe-12 followed by LC-MS-MS analysis. The abundance of CAT2, and its 2 isoforms, CAT1, and CAT3 are all higher than WT in both AtSAP5-oe-11 and AtSAP5-oe-12 (Table S3). This indicated that AtSAP5 enhances the CAT2 protein accumulation, but it not through transcriptional induction.

### Overexpression of AtSAP5 inhibits autophagosome formation in *Arabidopsis*

A previous report indicated that CAT2 is degraded through the ROS-induced autophagy pathway (*46*). Thus, we examined the role of AtSAP5 in autophagosome formation under H_2_O_2_ induction in WT (Col-0) and transgenic AtSAP5 overexpressing lines (AtSAP5-oe-11, AtSAP5-oe-12). We treated *Arabidopsis* with methyl viologen (MV), a known ROS inducer that can activate autophagy in plants (*47*). The fluorescence image of the roots revealed a substantial decrease in autophagosome formation in the AtSAP5 overexpression lines (Fig. 3D and 3E). This suggests that AtSAP5-mediated CAT2 protein accumulation is through the inhibition of autophagy-mediated CAT2 degradation.

### AtSAP5 does not directly enhance CAT2 activity but counteracts the negative effect of SA on CAT2 catalase activity

To analyze the effect of AtSAP5 on CAT2 activity, we performed *in vitro* CAT2 catalase activity assays using recombinant AtSAP5-His and CAT2-His proteins. CAT2-His activity was assayed *in vitro* with or without AtSAP5. The CAT2-His protein incubated with H_2_O_2_ showed higher CAT2 enzyme activity than the GFP-His control (1.0 to 0.03) (Fig. 4A, lanes 1 and 2). However, the CAT2 activities of CAT2-His protein alone (Fig. 4A, lane 2), CAT2-His with AtSAP5-His (Fig. 4A lanes 3–5), and with heat-inactivated AtSAP5-His protein (X-AtSAP5-His) (Fig. 4A lanes 6–8) were all similar. These results suggest that AtSAP5 neither enhances nor decreases CAT2 activity.

**Fig. 4.**
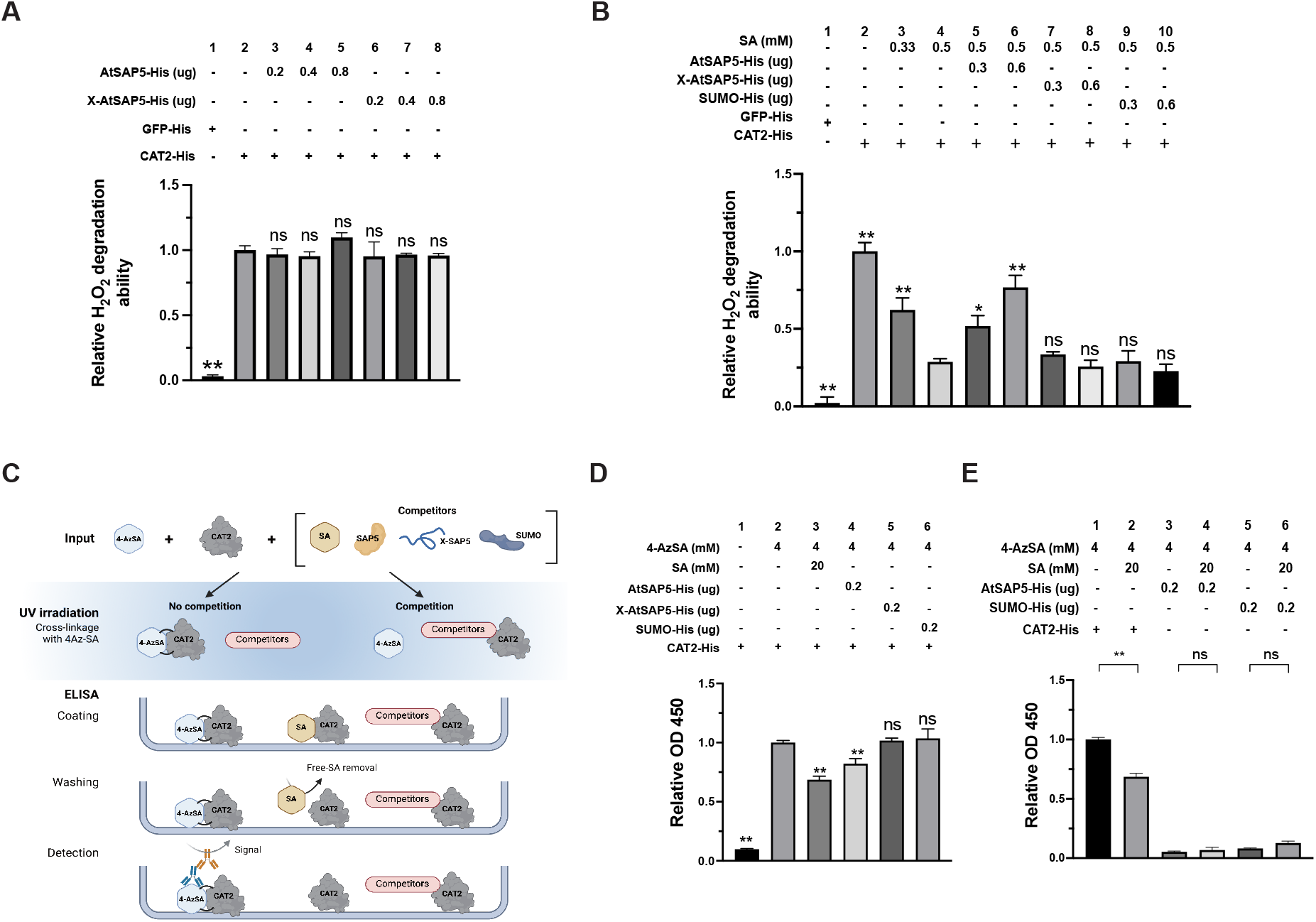
AtSAP5 has no effect on CAT2 catalase activity and does not bind to SA but counteracts the negative effect of salicylic acid (SA) on CAT2 activity through competition. The recombinant AtSAP5-His, heat-inactivated AtSAP5 (X-AtSAP5-His), GFP-His proteins were purified from *E coli*, and CAT2-His were purified from plants. **(A)** The *in vitro* H_2_O_2_ degradation ability of CAT2 was analyzed by incubating CAT2-His alone or mixed with AtSAP5-His or X-AtSAP5-His. GFP-His alone was used as a control. (**B)** The *in vitro* H_2_O_2_ degradation ability of CAT2 was analyzed by incubating CAT2-His mixed with AtSAP5-His, X-AtSAP5-His or SUMO-His in the presence or absence of SA. GFP-His and CAT2-His alone were used as controls. (**A** and **B**) The H_2_O_2_ degradation ability of CAT2-His alone was set to 1. Data represent mean ± SD; The results were combined from 3 independent experiments; **, *P* < 0.01; *, *P* < 0.05, Student’s *t*-test, compared with CAT2-His alone (**A**) or CAT2-His mixed with SA (0.5 mM) in (**B**). **(C)** Schematic representation of the ELISA assay. Recombinant CAT2-His mixed with the photoreactive SA analog 4-AzidoSA (4-AzSA) was incubated with SA, AtSAP5-His, X-AtSAP5, or SUMO-His for competition. Each reaction (input) was illuminated with UV light for crosslinking, then coated on a well in a 96 ELISA plate and detected by anti-SA antibody. The OD 450 value was measured from each reaction. **(D** and **E)** SA-binding competition assay was performed using photoaffinity labeling and ELISA. (**D**) CAT2-His mixed with the 4-AzSA was incubated with SA, AtSAP5-His, X-AtSAP5, or SUMO-His for the ELISA competition assay. CAT2-His alone with or without 4-AzSA were used as controls. (**E**) CAT2-His, AtSAP5-His, or SUMO-His mixed with 4-AzSA was incubated with or without SA for SA binding assay using ELISA. **(D** and **E)** The absorbance of CAT2-His protein incubated with 4-AzSA was set to 1. Data represent mean ± SD; The results were combined from 3 independent experiments **, *P* < 0.01, comparisons using Student’s *t*-test are indicated.

Since SA has been reported to inhibit *Arabidopsis* CAT2 activity (*25*), we next analyzed the effect of AtSAP5 on CAT2 enzyme activity in the presence of SA. The CAT2-His protein incubated with H_2_O_2_ showed much higher catalase activity than the GFP-His control (1.0 to 0.02) (Fig. 4B, lanes 1 and 2). Similar to previous reports (*19, 25*), the CAT2-His catalase activity decreased when SA was added (0.62 and 0.28) (Fig. 4B, lanes 3 and 4). However, in the presence of SA, increasing amounts of AtSAP5-His protein increased the CAT2 catalase activity (0.52 and 0.76) (Fig. 4B, lanes 5 and 6). This increase in CAT2 activity was not observed when heat-inactivated AtSAP5-His protein or the recombinant SUMO-His protein (with a similar molecular weight as AtSAP5-His) was used as a control (Fig. 4B, lanes 7–10). This indicates that AtSAP5 counteracts the negative effect of SA on CAT2 catalase activity *in vitro*.

These assays indicate that AtSAP5 does not enhance CAT2 activity but counteracts the negative effect of SA on CAT2.

### AtSAP5 interferes with SA-CAT2 binding

To analyze how AtSAP5 counteracts the negative effect of SA on CAT2, we performed a competition assay. A photoreactive SA analog, 4-AzidoSA (4-AzSA) was used in our assay (*48*). Under UV illumination, 4-AzSA can be cross-linked to its interacting protein, which allows the 4-AzSA-protein complex to be detected by anti-SA antibody. We designed an enzyme-linked immunosorbent assay (ELISA) using anti-SA antibodies to analyze whether *Arabidopsis* CAT2 can bind to SA and whether AtSAP5 can compete with the binding of SA to CAT2 (Fig. 4C).

We first incubated the purified CAT2-His proteins in the absence (Fig. 4D, lane 1) or presence of 4-AzSA (Fig. 4D, lane 2), and cross-linked them using UV irradiation. The result showed that a strong signal was detected in the reaction with CAT2-His plus 4-AzSA than CAT2-His alone (1 to 0.09) (Fig. 4D, lanes 1–2). Next, we included SA, AtSAP5-His, heat-inactivated AtSAP5-His (X-AtSAP5-His), or SUMO-His as competitors in the reaction (Fig. 4D, lanes 3–6). The result showed that SA decreased the 4-AzSA-CAT2-His signal (0.68) as expected (Fig. 4D, lane 3). In addition, AtSAP5-His (0.82) but not the control heat-inactivated AtSAP5-His (1.0) or SUMO-His (1.0) decreased the CAT2-His-4-AzSA signal (Fig. 4D, lanes 4–6). The results indicated that similar to SA, AtSAP5-His can prevent the interaction between 4-AzSA and CAT2-His.

We further analyzed whether AtSAP5-His can directly bind to SA, which will compete with CAT2-His to bind SA and decrease the CAT2-His-SA interaction. The purified CAT2-His, AtSAP5-His, or SUMO-His proteins mixed with 4-AzSA were incubated in the absence (Fig. 4E, lanes 1, 3 and 5) or presence of free SA (Fig. 4E, lanes 2, 4 and 6), and cross-linked using UV irradiation. The ELISA result showed that CAT2-His plus 4-AzSA showed the strongest signal (1.0), and AtSAP5-His (0.05) or SUMO-His (0.08) plus 4-AzSA only showed very weak signals (Fig. 4D, lanes 3-6). The results indicate that AtSAP5-His does not directly bind to SA.

Taken together, our results suggest that AtSAP5-His binds to CAT2-His and interferes with the interaction between SA and CAT2-His, which counteracts the negative effects of SA on CAT2-His activity.

## Discussion

Despite growing evidence indicating that SAPs play important roles in plant immunity, their regulatory mechanism has not been previously resolved. Most SAPs exhibit E3 ligase and ubiquitin-binding activity (*45*). Here, we found that, instead of acting as an E3 ligase or binding to ubiquitinated protein, AtSAP5 coordinates with SA and CAT2, which forms a delicate regulatory mode of action to fine-tune the H_2_O_2_. As SA-induced NPR1-dependent plant immunity activation largely depends on redox status (*12*), our findings provide a mechanistic explanation of how SA, CAT2, and AtSAP5 regulate redox status to promote the NPR1 nucleus translocation for activation of plant immunity.

We integrated our results into the current knowledge about SA-mediated plant immunity and have proposed a working model (Fig. 5). In the native state, the basal level of AtSAP5 and SA compete for binding to CAT2, and maintaining the homeostasis of H_2_O_2_ (Fig. 5). Pathogen infection induces SA accumulation (*49*). The accumulation of SA binds to CAT2, which inhibits CAT2 activity, resulting in an oxidative state, and serves as an initial signal for activation of plant immunity (*18, 19, 25*) (Fig. 5). The increased SA also upregulates the expression of AtSAP5 (*32*). Thereafter, more AtSAP5 binds to CAT2, thus counteracting the negative effect of SA on CAT2 activity (Fig. 5). The enhanced CAT2 activity further prevents the accumulation of H_2_O_2_ and H_2_O_2_-induced autophagosome formation, which forms a positive loop that further contributes to the accumulation of CAT2 (Fig. 5). Then, the cells shift from an oxidative state to a more reduced state. The reduced state promotes NPR1 monomer formation and translocates to the nucleus (*12*) where it activates downstream immune-response genes (Fig. 5).

**Fig. 5.**
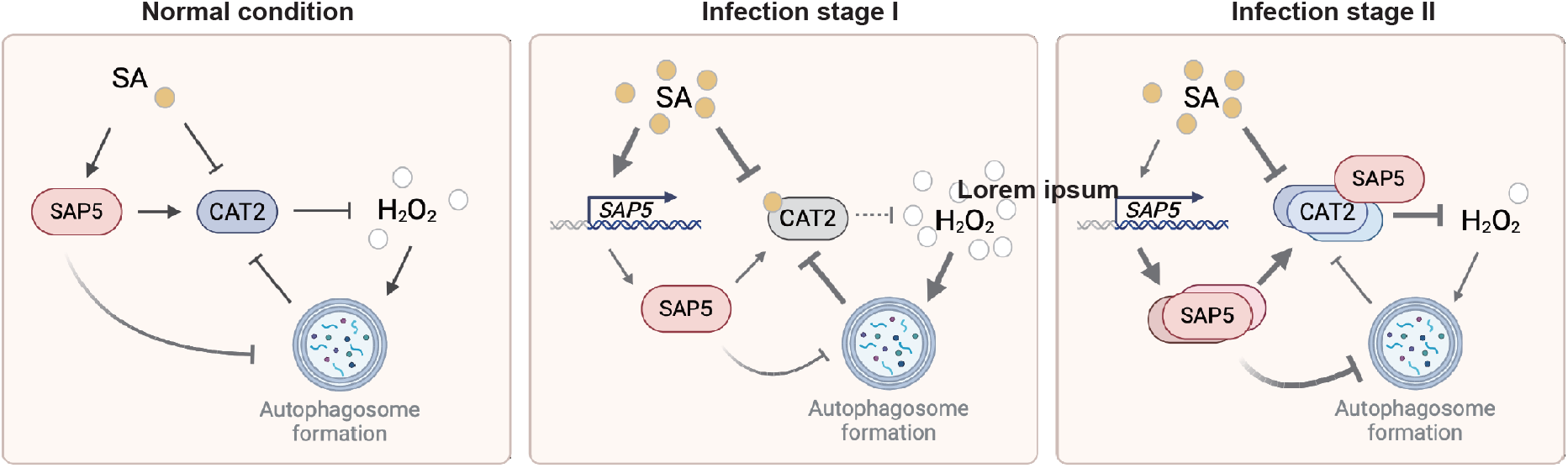
A model illustrating how AtSA5 transduces the SA signal to activate downstream immune-responsive genes. In the normal conditions, hydrogen peroxide (H_2_O_2_) is maintained in homeostasis. During pathogen infection stage I, the accumulated SA binds to CAT2 and inhibits CAT2 activity, leading to elevated H_2_O_2_ accumulation and an oxidative state in the cell. SA also activates the expression of AtSAP5. In the infection stage II, the accumulated AtSAP5 competes with SA for CAT2 binding, thereby protecting catalase activity from SA inhibition. This leads to a decrease in H_2_O_2_ levels and inhibits H_2_O_2_-induced autophagosome formation, resulting in the further accumulation of CAT2, which forms a positive feedback loop to promote additional CAT2 accumulation. (C)The enhanced CAT2 further decreases the H_2_O_2,_ leading to cell transit from the oxidative state to the relatively reduced state. Under a relatively reduced state, the oxidative NPR1 is reduced and enters the nucleus to activate downstream immune-responsive gene expression.

Similar to AtSAP5, human AWP1 and ZNF216 also contain A20-AN1 domains and participate in regulating the NF-kB-mediated immune pathway; however, the underlying mechanism is still unclear (*50, 51*). CATs are also conserved across kingdoms (*24, 52*). Whether human SAP-like proteins also have regulatory modes of action similar to those of AtSAP5 in plants will be an interesting subject for further study.

## Acknowledgments

We thank the assistance from Chuan-Chih Hsu at the Proteomics Core Lab, Institute of Plant and Microbial Biology, Academia Sinica; the assistance from Shu-Chen Shen for microscope imaging technical support at the Advanced Optical Microscope Core Facility, Agricultural Biotechnology Research Center at Academia Sinica. This work was supported by grants from Academia Sinica, Taipei, National Science and Technology Council, Taiwan (108-2313-B-001-010-MY3, 108-2811-B-001-583, 109-2811-B-001-509,110-2811-B-001-568) of Taiwan.

The funders had no role in the study design, data collection, and interpretation, or the decision to submit the work for publication.

## Funding

Academia Sinica

National Science and Technology Council (108-2313-B-001-010-MY3, 108-2811-B-001-583, 109-2811-B-001-509,110-2811-B-001-568)

## Author contributions

Conceptualization: LC, YZ, HHY

Methodology: LC, YSC, APC, SHC

Investigation: LC, APC, YSC, SHC

Visualization: LC, APC, YSC, SHC

Funding acquisition: HHY

Project administration: HHY

Supervision: HHY

Writing – original draft: LC, YZ, HHY

Writing – review & editing: LC, YZ, HHY

## References and Notes

1. D. F. Klessig, H. W. Choi, D. M. A. Dempsey, Systemic acquired resistance and salicylic acid: past, present, and future. Molecular plant-microbe interactions 31, 871–888 (2018).

2. Z. Q. Fu, X. Dong, Systemic acquired resistance: turning local infection into global defense. Annual review of plant biology 64, 839–863 (2013).

3. H. Cao, S. A. Bowling, A. S. Gordon, X. Dong, Characterization of an Arabidopsis mutant that is nonresponsive to inducers of systemic acquired resistance. The Plant Cell 6, 1583–1592 (1994).

4. T. Delaney, L. Friedrich, J. Ryals, Arabidopsis signal transduction mutant defective in chemically and biologically induced disease resistance. Proceedings of the National Academy of Sciences 92, 6602–6606 (1995).

5. J. Shah, F. Tsui, D. F. Klessig, Characterization of a s alicylic a cid-i nsensitive mutant (sai1) of Arabidopsis thaliana, identified in a selective screen utilizing the SA-inducible expression of the tms2 gene. Molecular Plant-Microbe Interactions 10, 69–78 (1997).

6. D. Wang, N. Amornsiripanitch, X. Dong, A genomic approach to identify regulatory nodes in the transcriptional network of systemic acquired resistance in plants. PLoS Pathogens 11, e123 (2006).

7. W. Wang et al., Structural basis of salicylic acid perception by Arabidopsis NPR proteins. Nature, 1–6 (2020).

8. Y. Ding et al., Opposite roles of salicylic acid receptors NPR1 and NPR3/NPR4 in transcriptional regulation of plant immunity. Cell 173, 1454–1467. e1415 (2018).

9. H. Cao, J. Glazebrook, J. D. Clarke, S. Volko, X. Dong, The Arabidopsis NPR1 gene that controls systemic acquired resistance encodes a novel protein containing ankyrin repeats. Cell 88, 57–63 (1997).

10. J. Ryals et al., The Arabidopsis NIM1 protein shows homology to the mammalian transcription factor inhibitor I kappa B. The Plant Cell 9, 425–439 (1997).

11. J. Withers, X. Dong, Posttranslational modifications of NPR1: a single protein playing multiple roles in plant immunity and physiology. PLoS pathogens 12, e1005707 (2016).

12. Z. Mou, W. Fan, X. Dong, Inducers of plant systemic acquired resistance regulate NPR1 function through redox changes. Cell 113, 935–944 (2003).

13. Y. Tada et al., Plant immunity requires conformational charges of NPR1 via S-nitrosylation and thioredoxins. Science 321, 952–956 (2008).

14. A. Saleh et al., Posttranslational modifications of the master transcriptional regulator NPR1 enable dynamic but tight control of plant immune responses. Cell Host & Microbe 18, 169–182 (2015).

15. A. M. Murphy, T. Zhou, J. P. Carr, An update on salicylic acid biosynthesis, its induction and potential exploitation by plant viruses. Current opinion in virology 42, 8–17 (2020).

16. J. Chen et al., NPR1 promotes its own and target gene expression in plant defense by recruiting CDK8. Plant physiology 181, 289–304 (2019).

17. A. Herrera-Vásquez, P. Salinas, L. Holuigue, Salicylic acid and reactive oxygen species interplay in the transcriptional control of defense genes expression. Frontiers in plant science 6, 171 (2015).

18. Z. Chen, H. Silva, D. F. Klessig, Active oxygen species in the induction of plant systemic acquired resistance by salicylic acid. Science 262, 1883–1886 (1993).

19. U. Conrath, Z. Chen, J. R. Ricigliano, D. F. Klessig, Two inducers of plant defense responses, 2, 6-dichloroisonicotinec acid and salicylic acid, inhibit catalase activity in tobacco. Proceedings of the National Academy of Sciences 92, 7143–7147 (1995).

20. J. Durner, D. F. Klessig, Salicylic acid is a modulator of tobacco and mammalian catalases. Journal of Biological Chemistry 271, 28492–28501 (1996).

21. D. Wendehenne, J. Durner, Z. Chen, D. F. Klessig, Benzothiadiazole, an inducer of plant defenses, inhibits catalase and ascorbate peroxidase. Phytochemistry 47, 651–657 (1998).

22. J. Durner, D. F. Klessig, Inhibition of ascorbate peroxidase by salicylic acid and 2, 6-dichloroisonicotinic acid, two inducers of plant defense responses. Proceedings of the National Academy of Sciences 92, 11312–11316 (1995).

23. M. Kvaratskhelia, S. J. George, R. N. Thorneley, Salicylic acid is a reducing substrate and not an effective inhibitor of ascorbate peroxidase. Journal of Biological Chemistry 272, 20998–21001 (1997).

24. A. Mhamdi et al., Catalase function in plants: a focus on Arabidopsis mutants as stress-mimic models. Journal of experimental botany 61, 4197–4220 (2010).

25. H.-M. Yuan, W.-C. Liu, Y.-T. Lu, CATALASE2 coordinates SA-mediated repression of both auxin accumulation and JA biosynthesis in plant defenses. Cell host & microbe 21, 143–155 (2017).

26. S. Vij, A. K. Tyagi, A20/AN1 zinc-finger domain-containing proteins in plants and animals represent common elements in stress response. Functional & integrative genomics 8, 301–307 (2008).

27. J. Giri et al., SAPs as novel regulators of abiotic stress response in plants. BioEssays 35, 639–648 (2013).

28. H. Tyagi, S. Jha, M. Sharma, J. Giri, A. K. Tyagi, Rice SAPs are responsive to multiple biotic stresses andoverexpression of OsSAP1, an A20/AN1 zinc-finger protein, enhancesthe basal resistance against pathogen infection in tobacco. Plant Science 225, 68–76 (2014).

29. S. Liu et al., Tomato stress-associated protein 4 contributes positively to immunity against necrotrophic fungus Botrytis cinerea. Molecular Plant-Microbe Interactions 32, 566–582 (2019).

30. M. Kang et al., Arabidopsis stress associated protein 9 mediates biotic and abiotic stress responsive ABA signaling via the proteasome pathway. Plant, cell & environment 40, 702–716 (2017).

31. S. Liu et al., Tomato SlSAP3, a member of the stress-associated protein family, is a positive regulator of immunity against Pseudomonas syringae pv. tomato DC3000. Molecular plant pathology 20, 815–830 (2019).

32. L. Chang et al., Plant A20/AN1 protein serves as the important hub to mediate antiviral immunity. PLoS pathogens 14, e1007288 (2018).

33. J. Zhao et al., The root-knot nematode effector MiPDI1 targets a stress-associated protein (SAP) to establish disease in Solanaceae and Arabidopsis. New Phytologist 228, 1417–1430 (2020).

34. L. Chang et al., Stress associated proteins coordinate the activation of comprehensive antiviral immunity in Phalaenopsis orchids. New Phytologist 233, 145–155 (2022).

35. L. Chang et al., Plant A20/AN1 proteins coordinate different immune responses including RNAi pathway for antiviral immunity. bioRxiv, 622696 (2019).

36. H. Tyagi, S. Jha, M. Sharma, J. Giri, A. K. Tyagi, Rice SAPs are responsive to multiple biotic stresses and overexpression of OsSAP1, an A20/AN1 zinc-finger protein, enhances the basal resistance against pathogen infection in tobacco. Plant Science 225, 68–76 (2014).

37. H. Cao, S. A. Bowling, A. S. Gordon, X. Dong, Characterization of an Arabidopsis mutant that is nonresponsive to inducers of systemic acquired resistance. The Plant Cell 6, 1583–1592 (1994).

38. J. D. Clarke, Y. Liu, D. F. Klessig, X. Dong, Uncoupling PR gene expression from NPR1 and bacterial resistance: characterization of the dominant Arabidopsis cpr6-1 mutant. The Plant Cell 10, 557–569 (1998).

39. G. Felix, J. D. Duran, S. Volko, T. Boller, Plants have a sensitive perception system for the most conserved domain of bacterial flagellin. The Plant Journal 18, 265–276 (1999).

40. T. Boller, G. Felix, A renaissance of elicitors: Perception of microbe-associated molecular patterns and danger signals by pattern-recognition receptors. Annual Review of Plant Biology 60, 379–406 (2009).

41. S. Peleg-Grossman, N. Melamed-Book, G. Cohen, A. Levine, Cytoplasmic H2O2 prevents translocation of NPR1 to the nucleus and inhibits the induction of PR genes in Arabidopsis. Plant Signaling & Behavior 5, 1401–1406 (2010).

42. M. Zhang et al., Two cytoplasmic effectors of Phytophthora sojae regulate plant cell death via interactions with plant catalases. Plant physiology 167, 164–175 (2015).

43. S. Chaouch et al., Peroxisomal hydrogen peroxide is coupled to biotic defense responses by ISOCHORISMATE SYNTHASE1 in a daylength-related manner. Plant Physiology 153, 1692–1705 (2010).

44. M. Kang et al., At MBP-1, an alternative translation product of LOS 2, affects abscisic acid responses and is modulated by the E 3 ubiquitin ligase A t SAP 5. The Plant Journal 76, 481–493 (2013).

45. M. Kang, M. Fokar, H. Abdelmageed, R. D. Allen, Arabidopsis SAP5 functions as a positive regulator of stress responses and exhibits E3 ubiquitin ligase activity. Plant Molecular Biology 75, 451–466 (2011).

46. T. Hackenberg et al., Catalase and NO CATALASE ACTIVITY1 promote autophagy-dependent cell death in Arabidopsis. The Plant Cell 25, 4616–4626 (2013).

47. S. Han et al., Cytoplastic glyceraldehyde-3-phosphate dehydrogenases interact with ATG3 to negatively regulate autophagy and immunity in Nicotiana benthamiana. The Plant Cell 27, 1316–1331 (2015).

48. M. Tian et al., The combined use of photoaffinity labeling and surface plasmon resonance-based technology identifies multiple salicylic acid-binding proteins. The Plant Journal 72, 1027–1038 (2012).

49. T. E. Mishina, J. Zeier, Pathogen-associated molecular pattern recognition rather than development of tissue necrosis contributes to bacterial induction of systemic acquired resistance in Arabidopsis. The Plant Journal 50, 500–513 (2007).

50. J. Huang et al., ZNF216 is an A20-like and IκB kinase γ-interacting inhibitor of NFκB activation. Journal of Biological Chemistry 279, 16847–16853 (2004).

51. E.-J. Chang, J. Ha, S.-S. Kang, Z. H. Lee, H.-H. Kim, AWP1 binds to tumor necrosis factor receptor-associated factor 2 (TRAF2) and is involved in TRAF2-mediated nuclear factor-kappaB signaling. The international journal of biochemistry & cell biology 43, 1612–1620 (2011).

52. J. Switala, P. C. Loewen, Diversity of properties among catalases. Archives of biochemistry and biophysics 401, 145–154 (2002).

