## Supplemental Material for "Delicate tuning of H_2_O_2_ by salicylic acid, catalase 2, and AtSAP5 incites plant immunity"

**The supplementary materials includes:**

Materials and Methods
Figs. S1 to S2
Tables S1 to S4
References

**Materials and Methods**

**Plants**

The wild-type (WT) *Arabidopsis* (Col-0) and all transgenic *Arabidopsis* lines were maintained in a greenhouse with a controlled 12-h photoperiod (200 μ mol $m^{-2}s^{-2}$) at 22°C/22°C (day/night) for 3 to 4 weeks before analysis.

**Bacteria**

*Escherichia coli* strain BL21 was grown on Luria broth (LB) agar plates or in LB broth. *Pseudomonas syringae* pv. *tomato* DC30000 (*Pst* DC3000) was grown in King's B medium (20 g/L proteose peptone, 1.5 g/L $K_2HPO4$, 10 mL glycerol, and 1.5 g/L MgSO4 •$7H_2O$, pH 7.0).

**Transgenic *Arabidopsis***

The approach used to generate transgenic *Arabidopsis* was as described previously (*1*) with some modification. For the generation of transgenic *Arabidopsis* lines overexpressing AtSAP5, the plants were transformed using the floral dip method with *Agrobacterium tumefaciens* strain GV3101 carrying pAtSAP5-oe (*1*). The *Arabidopsis npr1-1* mutant (*2*) was transformed using the floral dip method with *A. tumefaciens* strain GV3101 carrying pNPR1-GFP and pAtSAP5-oe (*1*) to generate the transgenic plants *35S::NPR1-GFP* in *npr1-1* (NPR1-GFP/*npr1*) and 35S::AtSAP5 in *npr1-1* (AtSAP5-oe/*npr1*-1 and AtSAP5-oe/*npr1*-2), respectively.

***Pst* DC3000 inoculation and flagellin 22 (flg22) treatment**

For *Pst* DC3000 inoculation, leaves of 4-week-old *Arabidopsis* plants were infiltrated with *Pst* DC3000 (OD 0.0005). At 3 days post-inoculation, the inoculated leaves were ground in sterile water. Serial dilution was performed, and King's B medium containing 100 μg/ml rifampicin was used for colony counting.

For flg22 treatment, three leaves from 4-week-old *Arabidopsis* plants were infiltrated with 10 μM flg22 (Genscript Biotech Corporation, Piscataway, NJ, USA) using a syringe without a needle. The infiltrated leaves were collected at 24 h post-infiltration.

**RNA isolation and reverse transcription quantitative PCR (RT-qPCR)**

Total RNA extraction and RT-qPCR detection were performed as described previously (*1*), and the primer pairs are listed in Table S4.

**NPR1-GFP nucleus translocation assay**

For the construction of GFP-tagged NPR1, an RT-PCR reaction was performed to amplify the *NPR1* fragment (without a stop codon) using the primer pairs, NPR1-ORF-F/NPR1-ORF-non-stop-R (Table S4), with total RNA extracted from *Arabidopsis* (Col-0) as a template. The amplified fragment was cloned into the Gateway entry vector pENTR/D-TOPO (Thermo Fisher-Scientific, Waltham, MA, USA) to generate pENTR-NPR1-non-stop following the manufacturer's protocol. Subsequently, an LR Gateway cloning reaction (Thermo Fisher-Scientific) was performed to transfer the *NPR1* fragment from pENTR-NPR1-non-stop into pK7FWG2 to generate pNPR1-GFP with GFP at the C terminus of NPR1. The construction of the vector used for HA-tagged AtSAP5 overexpressing, pAtSAP5-oe, was previously described (*1*).

The approach for isolation of protoplasts from the transgenic *Arabidopsis* NPR1-GFP/*npr1* is described in the "**Transgenic *Arabidopsis***" section above, and transfection of pAtSAP5-oe was performed as previously described (*3*). Sixteen hours post transfection, the cells were treated with $H_2O$ (mock) or SA (1 mM) for 3 h, followed by the detection of fluorescence signals using confocal microscopy (Zeiss LSM 780, plus ELYRA S.1).

**Immunoprecipitation**

For identification of AtSAP5 interacting proteins, total proteins were extracted from leaves (0.2 g) of WT (Col-0) and transgenic *Arabidopsis* lines overexpressing AtSAP5 (AtSAP5-oe-11 and AtSAP5-oe-12) (*1*) with the use of 200 μl immunoprecipitation buffer (50 mM Tris-pH 7.5, 0.1% NP40, 10 mM $MgCl_2$, 150 mM NaCl, 10 μM MG132) and 1X complete protease inhibitor (Roche, Basel, Switzerland). The extract was centrifuged at 12,000 × g for 10 min at 4°C. Then, the supernatants were transferred to a non-stick tube and pre-cleared with protein A sepharose beads (GE Healthcare Life Sciences, Pittsburgh, PA, USA) for 1 h at 4°C with gentle shaking. Then, the extract was centrifuged at 1500 × g for 2 min at 4°C. The supernatant was further incubated with the anti-AtSAP5 antibody conjugated with sepharose beads (GE Healthcare Life Sciences) for 4 h at 4°C with gentle shaking. The beads were washed three times with ice-cold immunoprecipitation buffer and eluted with sodium dodecyl sulfate-polyacrylamide gel electrophoresis (SDS-PAGE) sample buffer. The eluted proteins were separated by SDS-PAGE and stained using a silver stain kit (Thermo Fisher-Scientific), followed by liquid chromatography tandem mass spectrometry (LC-MS/MS) for identification of the putative AtSAP5-interacting proteins. Candidate proteins identified in WT, AtSAP5-oe-11 and AtSAP5-oe-12 are listed in Table S1. The number of PSM for each candidate protein in WT, AtSAP5-oe-11, and AtSAP5-oe-12 is derived from three independent experiments.

For analysis of the ubiquitination status of CAT2, total proteins extracted from leaves (0.2 g) of WT (Col-0) and AtSAP5-oe-11 and AtSAP5-oe-12 were used for immunoprecipitation with anti-CAT antibody (Agrisera, Vännäs, Sweden) following the same approach described above. The identified proteins and the ubiquitination site are listed in Table S2.

**In-gel trypsin digestion**

The protocol used for in-gel trypsin digestion of proteins was adapted mainly from the method described by Wilm and Mann (*4*). Briefly, the protein bands from the gel were manually excised, and each band was cut into small pieces (~0.5 $mm^3$). The gel pieces in an Eppendorf tube were washed a few times with a solution containing 50% methanol and 5% acetic acid for 2–3 h. The gel pieces were washed with a solution of 25 mM $NH_4$ $HCO_3$ in 50% acetonitrile for 10 min twice. Then, the gel pieces were dried in a vacuum centrifuge. Proteins in the gel pieces were reduced with DTT and alkylated with iodoacetamide, then washed and dried in a vacuum centrifuge before trypsin digestion. The gel pieces were then incubated in 25–40 µL trypsin solution, 25 mM $NH_4HCO_3$ containing 75 to 100 ng of sequencing grade modified trypsin (Promega), for 12–16 h at 37°C. To recover the tryptic peptides, 30 µL of a solution of 5% formic acid and 50% acetonitrile was added to the gel pieces, which were agitated in a vortex for 30–60 min and collected into a new tube. The recovery process was repeated once with 15 µL solution, and the resulting recovered solutions were combined and dried in a vacuum centrifuge. The dried pellet was re-dissolved in 10–20 µL of 0.1% formic acid for LC-MS/MS analysis.

**MS method for protein identification and analysis**

An LC-nESI-Q Exactive mass spectrometer (Thermo Fisher Scientific) coupled with an online nanoUHPLC (Dionex UltiMate 3000 Binary RSLCnano) was used for protein analysis and identification. An Acclaim PepMap 100 C18 trap column (75 µm × 2.0 cm, 3 µm, 100 Å, Thermo

Fisher-Scientific) and an Acclaim PepMap RSLC C18 nano LC column (75 µm × 15 cm, 2 µm, 100 Å) were used to deliver solvent and separate tryptic peptides with a linear gradient from 3% to 30% of acetonitrile in 0.1% (v/v) formic acid for 3 h at a flow rate of 300 nL/min. The acquisition of the MS data was performed in the data-dependent mode with a full MS scan followed by 10 MS/MS scans of the top 10 precursor ions from the MS scan. The MS scan was performed with a resolving power of 70,000 over the mass-to-charge (m/z) range 380 to 1800 and dynamic exclusion enabled. The data-dependent MS/MS acquisitions were performed with a 2 m/z isolation window, 27 NCE, and 17,500 resolving power. Peptide and protein identification was performed by using the Proteome Discoverer software (Thermo Fisher Scientific) with the SEQUEST search engine to search against the *Arabidopsis* database (TAIR database, https://www.arabidopsis.org/). The parameters for database searches were set as follows: full trypsin digestion with 2 maximum missed cleavage sites, precursor mass tolerance = 10 ppm, fragment mass tolerance = 0.02 Da, dynamic modifications: oxidation (M), static modifications: carbamidomethyl (C). The identified peptides were validated by performing searches against the decoy database using the Percolator algorithm, which rescored peptide spectrum matches using q-values and posterior error probabilities. All the peptides were filtered with a q-value threshold of 0.01 (1% false discovery rate), and proteins were filtered with a minimum of two peptides per protein, in which only the rank 1 peptide and peptides in the top-scoring proteins were counted.

**Preparation and transfection of protoplasts for bimolecular fluorescence complementation assays.**

For the construction of AtSAP5 vectors used for bimolecular fluorescence complementation (BiFC) analysis, PCR was carried out to amplify the *AtSAP5* ORF (without a stop codon) using the primer pair AtSAP5-ORF-F/AtSAP5-ORF-non-stop-R (Table S4) with pAtSAP5-oe (*1*) as a template. The amplified fragment was cloned into the Gateway entry vector pENTR/D-TOPO (Thermo Fisher-Scientific) to generate pENTR-AtSAP5-non-stop following the manufacturer's protocol. Subsequently, an LR Gateway cloning reaction (Thermo Fisher Scientific) was performed to transfer the ORF fragment of *AtSAP5* from pENTR-AtSAP5-non-stop into pGWcY (NCBI GenBank: AB626695) to generate pAtSAP5-cY with AtSAP5 fused at the C-terminal end with the C terminus of YFP (cYFP). For the construction of CAT2 vectors used for BiFC analysis, total RNA extracted from *Arabidopsis* (Col-0) was used as a template to amplify the *CAT2* fragment through RT-PCR with the primer pair CAT2-ORF-F/CAT2-ORF-R (Table S4). The amplified fragments were cloned into the Gateway entry vector pENTR/D-TOPO (Thermo Fisher Scientific) to generate pENTR- pENTR-CAT2. Subsequently, an LR Gateway cloning reaction (Thermo Fisher Scientific) was performed to transfer the *CAT2* fragment into pnYGW (NCBI GenBank: AB626694) to generate pnY-CAT2 with the N terminus of CAT2 fused with the N terminus of YFP (nYFP).

*Arabidopsis* protoplast isolation and transfection were performed as previously described (*3, 5*). Fluorescence signals in transformed protoplasts were detected by confocal microscopy (Zeiss LSM 780, plus ELYRA S.1).

**Split luciferase assay**

For the construction of vectors used for the split luciferase assay, a PCR reaction was carried out to amplify *AtSAP5* and *CAT2* using the primer pairs BamH1-AtSAP5-F/SalI-AtSAP5-R (for *AtSAP5*) and KpnI-CAT2-F/SalI-CAT2-R (for *CAT2*) (Table S4), and pAtSAP5-oe (for *AtSAP5*) (*1*) and pENTR-CAT2 (for *CAT2*) were used as templates. The PCR-amplified fragments of *AtSAP5* were digested with BamH1 and SalI (NEB, Ipswich, MA, USA) and ligated into BamH1- and SalI-digested pCAMBIA1300-NLuc (TAIR stock centers, CD3-1699) to generate pAtSAP5-nLuc. The PCR-amplified fragments of *CAT2* were digested by KpnI and SalI (NEB) and ligated into KpnI- and SalI-digested pCAMBIA1300-CLuc (TAIR stock centers, CD3-1700) to generate pcLuc-CAT2.

The plasmids pAtSAP5-nLuc, pcLuc-CAT2, and pCAMBIA1300-CLuc pCAMBIA1300-CLuc were individually transformed into *Agrobacterium.* Co-expression experiments were performed by mixing two *Agrobacterium* strains carrying an individual construct in a 1:1 ratio, then infiltrating them into *N. benthamiana* leaves through agroinfiltration. Three days after inoculation, 1 mM luciferin was sprayed onto the inoculated leaves and chemiluminescence images were taken by a CCD camera.

**Construction of recombinant protein expression vectors**

For construction of the His-tagged CAT2 transient overexpression vector, plant total RNA extracted from *Arabidopsis* (Col-0) was used as a template to amplify *CAT2* by RT-PCR with the primer pair CAT2-ORF-F/CAT2-ORF-non-stop-R (Table S4). The amplified fragment was cloned into the Gateway entry vector pENTR/D-TOPO (Thermo Fisher-Scientific) to generate pENTR-CAT2-non-stop. Subsequently, an LR Gateway cloning reaction (Thermo Fisher-Scientific) was performed to transfer the *CAT2* fragment from pENTR-CAT2-non-stop into the 35S promoter-driven pGWB408 (*6*), to obtain pCAT2-His. The method for construction of pENTR-GFP-non-stop and His-tagged GFP transient overexpression vector, pGFP-His, was similar to that described above, except that the primer pair GFP-ORF-F/GFP-ORF-non-stop-R (Table S4) and template pGFP were used to generate the *GFP* fragment.

For construction of the His-tagged SUMO vector, full-length *sumo* was amplified by PCR with the primer pair NdeI-SUMO-F/NotI-SUMO-R (Table S4) with the pET His6 Sumo TEV LIC cloning vector (Addgene number: 29659) as a template. PCR-amplified gene fragments were cloned into the pET24b expression vector (Novagen) with a C-terminal histidine tag (His-tag) to generate pET-sumo.

**Expression and purification of recombinant protein**

The agroinfiltration approach was used for expression of CAT2-His or GFP-His in *N. benthamiana* following a previously described method with some modification (*1*). Briefly, *A. tumefaciens* C58C1 (pTiB6S3ΔT)$^H$ competent cells were transformed with pCAT2-His, pGFP, or pBin61-p19 using an electroporation system (Bio-Rad Laboratories, Hercules, CA, USA). Then, the *A. tumefaciens* strains were incubated at 28°C until the optical density at 600 nm, $OD_{600}$, reached 0.8–1.0. After centrifugation, cells were resuspended in 20 mL AB-MES medium (17.2 mM $K_2HPO_4$, 8.3 mM $NaH_2PO_4$, 18.7 mM $NH_4Cl$, 2 mM KCl, 1.25 mM $MgSO_4$, 100 µM $CaCl_2$, 10 µM $FeSO_4$, 50 mM MES, 2% glucose [$wv^{-1}$], pH 5.5) with 200 µm acetosyringone (*7*) and cultured overnight. The overnight culture was centrifuged (3000 × g, 10 min, at room temperature), the supernatant was removed, and the culture was resuspended in infiltration medium (20 mL) containing 50% Murashige and Skoog medium (1/2 MS salts supplemented with 0.5% sucrose (w $v^{-1}$), pH 5.5), 50% AB-MES, and 200 µm acetosyringone (*7*). The infiltration medium containing the transformed *A. tumefaciens* was used for infiltration.

For expression and purification of CAT2-His or GFP-His in *N. benthamiana*, the *Agrobacterium* carrying pCAT2-His or pGFP-His mixed with pBin61-p19 in a 1:1 ratio was co-infiltrated into *N. benthamiana* leaves at $OD_{600}$=1. Three days post infiltration, total proteins from the infiltrated region (25 g) were extracted with 50 mL immunoprecipitation buffer (50 mM Tris-pH 7.5, 0.1% NP40, 10 mM $MgCl_2$, 150 mM NaCl, 10 μM MG132) with 1× complete protease inhibitor (Roche). The extract was incubated at 4°C for 30 min with shaking (30 rpm), followed by sonication for 1 min. Then, the extracts were centrifuged at 10,000 rpm for 10 min at 4°C, and the supernatants were passed through a filter (44 μm cell strainer) and collected in a new tube. After centrifugation at 10,000 rpm for 10 min at 4°C, the supernatants were collected in a new tube. Then the His-tagged recombinant proteins were purified by TALON Super-flow (GE Healthcare Life Sciences) according to the manufacturer's instructions. The elution was carried out with 250 mM imidazole.

The approach for purification of SUMO-His and AtSAP5-His recombinant proteins from *E. coli* was as previously described (*1*).

***In vitro* pull-down assays**

For the AtSAP5 pull-down assay, the recombinant AtSAP5-His, GFP-His, and CAT2-His proteins were purified as described in the section "**Expression and purification of recombinant protein**". Each reaction was performed in 500 μL immunoprecipitation buffer as described in the section **"Immunoprecipitation"** with recombinant AtSAP5-His, GFP-His, or CAT2-His. The reactions were incubated for 2 h with gentle shaking at 4°C. Then, the reactions were incubated with anti-AtSAP5 antibody and protein A Sepharose beads (GE Healthcare Life Sciences) for 4 h with gentle shaking at 4°C. Afterwards, the beads were washed three times with ice-cold immunoprecipitation buffer and eluted with SDS-PAGE sample buffer. The eluted proteins were separated by SDS-PAGE and detected by immunoblotting using the anti-His antibody (Applied Biological Materials, Richmond, Canada).

***In vitro* ubiquitination assay**

*In vitro* ubiquitination assays were performed as described (*1*) with modification. An amount of 3 μg purified His-tagged AtSAP5-His, and CAT2-His recombinant proteins were used for each ubiquitination reaction. Reactions were incubated at 30°C for 3 hours and analyzed by SDS-PAGE followed by immunoblot analysis using anti-FLAG antibodies (Sigma) or anti-CAT antibody (Agrisera).

**$H_2O_2$-scavenging activity assay**

For the $H_2O_2$-scavenging activity assay in *Arabidopsis*, leaves (0.1 g) from the WT (Col-0), transgenic *Arabidopsis* lines overexpressing AtSAP5 (AtSAP5-oe-11, and AtSAP5-oe-12) were collected, and the CAT activity was analyzed using a previously described approach (*8*). Total protein from collected leaf samples was extracted using extraction buffer (50 mM phosphate buffer, pH 7.0, 1% TritonX-100) with 1× complete protease inhibitor (Roche). Then, the extracted protein was added into 300 μL buffered substrate (50 mM phosphate buffer, pH 7.0, 10 mM $H_2O_2$). The decomposition of $H_2O_2$ was followed by measuring the decline in absorbance at 240 nm. The change of 240 nm values per min represented the CAT2 activity.

**Methyl viologen (MV) treatment and assay**

Seven-day-old seedlings grown in ½ MS solid media were transferred to the same media containing 10 uM MV for the treatment (*9*). Two days after the treatment, MV-treated seedlings were transferred to a new 6-well plate and stained with 2 ml of 50 uM Monodansylcadeverine (MDC) for 10 min in the dark, followed by two washes using 1X Phosphate-buffered saline (PBS) buffer (*10*). MDC signals were visualized using DAPI filter, excited at 325-375 nm and emission collected at 435-485 nm using a Carl Zeiss AxioImager Z1 fluorescence microscope.

**In vitro CAT2 enzyme activity assay**

The *in vitro* CAT2 activity assay was as described previously (*8*). In brief, CAT2-His protein (2 μg) (purified from *N. benthamiana*) alone or mixed with different amounts of SA, AtSAP5-His, heat-inactivated AtSAP5 (X-AtSAP5-His), or SUMO-His was added to 300 μL buffered substrate (50 mM phosphate buffer [pH 7.0] and 10 mM $H_2O_2$). The decomposition of $H_2O_2$ was followed by measuring the decline in absorbance at 240 nm. The change in 240 nm values per min were used to represent the CAT2 activity. The recombinant GFP-His protein was used as a control.

**SA-binding activity determined by photoaffinity labeling and enzyme-linked immunosorbent assay**

SA-binding activity was assessed by photoaffinity labeling as described previously (*11*). For Fig. 4D, the purified CAT2-His recombinant protein alone or mixed with SA, AtSAP5-His, heat-inactivated AtSAP5 (X-AtSAP5-His), or SUMO-His was incubated for 1 h on ice with 4-AzidoSA

(4-AzSA) (4 mM) in 1× PBS (50 µL) followed by illumination with 254 nm UV light at an energy level of 30 mJ. For Fig 4E, the purified CAT2-His, AtSAP5-His, or SUMO-His recombinant protein alone or mixed with SA was incubated for 1 h on ice with 4-AzSA (4 mM) in 1× PBS (50 µL) followed by illumination with 254 nm UV light at an energy level of 30 mJ. Then, the reaction mixture (20 µL) was used for an enzyme-linked immunosorbent assay (ELISA) with anti-SA antibody (GeneTex, Irvine, CA, USA) to detect 4-AzSA-crosslinked proteins.

For the ELISA assay, the reaction mixture (20 µL) described above was incubated with 200 µL of coating buffer (1× PBS) at 37°C for 1 h. After incubation, the plate was washed three times with PBST (phosphate-buffered saline with 0.05% Tween-20) followed by blocking using 1% BSA (dissolved in 1× PBS) at room temperature for 1 h. Then, the blocking reagent was discarded, followed by incubation of 200 µL conjugate buffer (1× PBS with 1% BSA) containing anti-SA antibody (GeneTex) at 4°C overnight. After washing three times using PBST, the plate was incubated with 200 µL conjugate buffer (1× PBS with 1% BSA) containing anti-rabbit antibody (Thermo Fisher Scientific) at 37°C for 1 h. After washing three times using PBST, the plate was incubated with 100 µL TMB (SeraCare, Milford, MA, USA) for development following the manufacturer's protocol. Then, the plate was analyzed by measurement of absorbance at 450 nm.

### Quantification and Statistical Analysis

The statistical analyses were performed in GraphPad Prism version 9.0. Data are presented as mean ± SD. The pair-wise Student's *t*-test was performed to analyze the statistical significance of differences between samples.

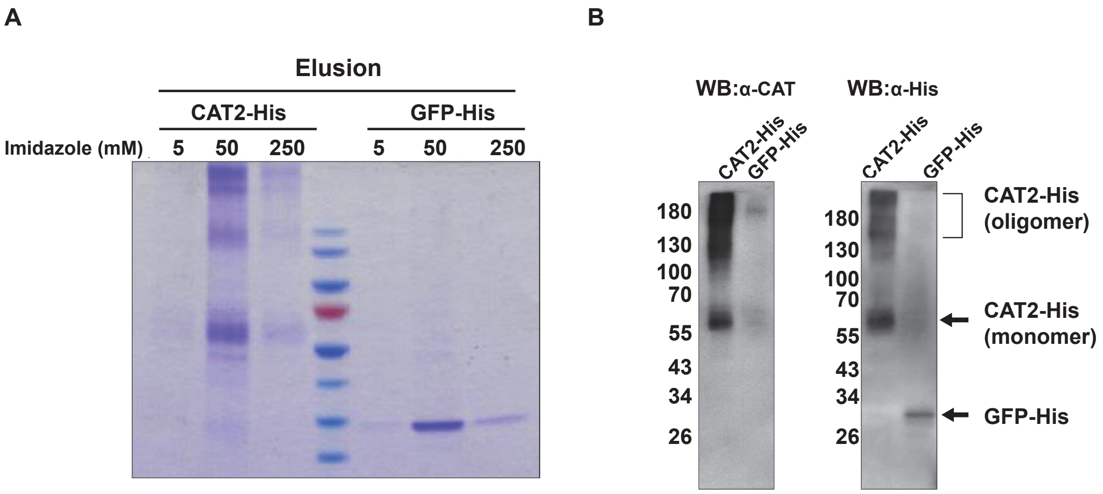


**Fig. S1 Recombinant CAT2-His and GFP-His proteins purified from *Nicotiana benthamiana.***
**(A)** The recombinant CAT2-His and GFP-His proteins purified from *N. benthamiana* were separated by SDS-PAGE and stained with Coomassie blue. Different concentrations of imidazole used for elusion are indicated. **(B)** Two sets of CAT2-His and GFP-His proteins from (A) were separated by SDS-PAGE and detected with anti-CAT2 or anti-His antibody. The oligomer and monomer forms of recombinant CAT2-His and GFP-His proteins are shown. The CAT2-His monomer and oligomer were both detected, as observed in a previous study (*12*).

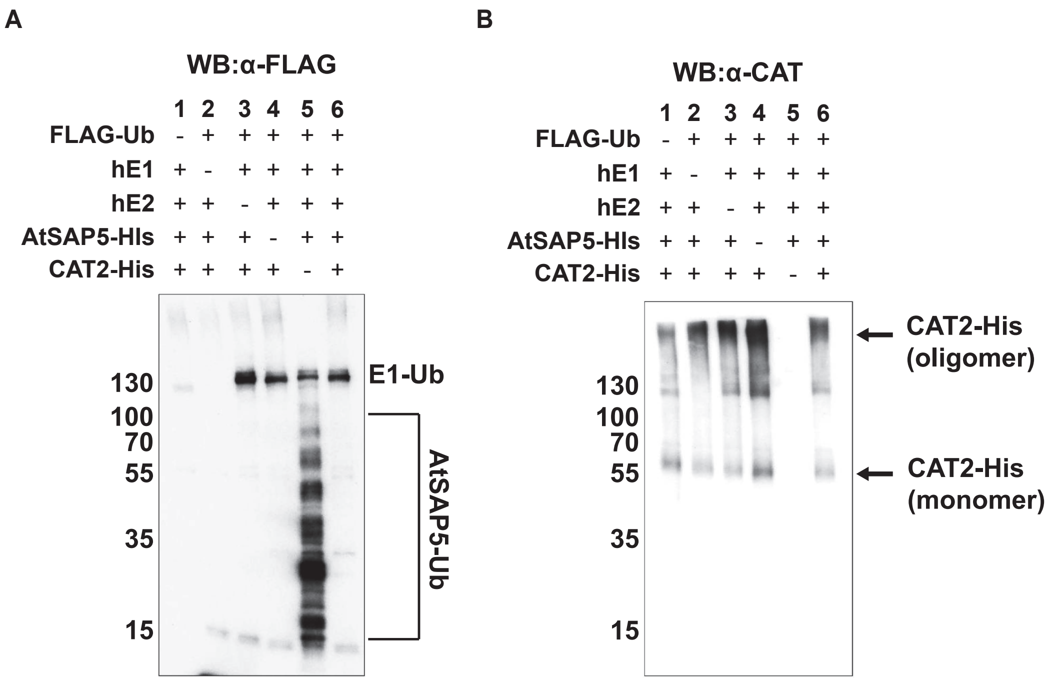


**Fig. S2** ***In vitro*** **ubiquitination assay. (A and B)** An *in vitro* ubiquitination assay was performed by incubating different combinations of the following proteins: FLAG-Ub, human ubiquitin-activating enzyme (hE1), human ubiquitin-conjugating enzyme (hE2), AtSAP5-His, and CAT2-His. Proteins added (+) or omitted (-) in each assay are indicated. Ubiquitinated proteins were analyzed by immunoblotting using **(A)** anti-FLAG antibody or **(B)** anti-CAT2 antibody. Ubiquitinated AtSAP5 (AtSAP5-Ub), E1 conjugated with Ub (E1-Ub), and CAT2-His (oligomer and monomer) are indicated.

**Table S1** AtSAP5 interacting proteins identified in WT, AtSAP5-oe-11 and AtSAP5-oe-12 by co-immunoprecipitation

| **ID**[a] | **Alias** | **PSM**[b] | | |
|---|---|---|---|---|
| | | WT | AtSAP5-oe-11 | AtSAP5-oe-12 |
| AT2G42600.2 | phosphoenolpyruvate carboxylase 2 | 214 | 237 | 239 |
| AT1G53310.3 | phosphoenolpyruvate carboxylase 1 | 137 | 167 | 161 |
| ATCG00480.1 | ATP synthase subunit beta | 92 | 112 | 85 |
| AT5G02940.1 | protein of unknown function (DUF1012) | 72 | 76 | 75 |
| ATCG00490.1 | ribulose-bisphosphate carboxylases | 72 | 95 | 94 |
| AT2G14120.3 | dynamin related protein | 68 | 59 | 67 |
| AT2G42490.1 | Copper amine oxidase family protein | 66 | 78 | 67 |
| ATCG00120.1 | ATP synthase subunit alpha | 58 | 54 | 51 |
| AT4G33650.2 | dynamin-related protein 3A | 40 | 42 | 42 |
| AT5G22800.1 | Alanyl-tRNA synthetase, class IIc | 38 | 54 | 56 |
| AT1G20620.1 | catalase 3 | 33 | 41 | 43 |
| AT3G26650.1 | glyceraldehyde 3-phosphate dehydrogenase A subunit | 33 | 33 | 29 |
| AT4G35090.1 | catalase 2 | 32 | 42 | 39 |
| AT3G12780.1 | phosphoglycerate kinase 1 | 31 | 28 | 22 |
| AT1G42970.1 | glyceraldehyde-3-phosphate dehydrogenase B subunit | 30 | 29 | 35 |
| AT4G04640.1 | ATPase, F1 complex, gamma subunit protein | 27 | 29 | 17 |
| AT4G13670.1 | plastid transcriptionally active 5 | 25 | 51 | 30 |
| AT4G38970.1 | fructose-bisphosphate aldolase 2 | 24 | 18 | 25 |
| AT1G13440.1 | glyceraldehyde-3-phosphate dehydrogenase C2 | 22 | 29 | 18 |
| AT3G11630.1 | Thioredoxin superfamily protein | 21 | 20 | 20 |

| | | | | |
|---|---|---|---|---|
| AT5G06290.1 | 2-cysteine peroxiredoxin B | 20 | 22 | 17 |
| ATCG00350.1 | Photosystem I, PsaA/PsaB protein | 20 | 19 | 7 |
| AT2G39730.1 | rubisco activase | 19 | 18 | 20 |
| AT2G43180.1 | Phosphoenolpyruvate carboxylase family protein | 19 | 30 | 28 |
| AT2G13360.2 | alanine:glyoxylate aminotransferase | 16 | 25 | 29 |
| ATCG00340.1 | Photosystem I, PsaA/PsaB protein | 15 | 17 | 8 |
| AT1G07940.1 | GTP binding Elongation factor Tu family protein | 12 | 11 | 16 |
| AT3G02660.1 | Tyrosyl-tRNA synthetase, class Ib, bacterial/mitochondrial | 11 | 23 | 15 |
| AT5G38410.3 | Ribulose bisphosphate carboxylase (small chain) family protein | 11 | 14 | 8 |
| ATCG00280.1 | photosystem II reaction center protein C | 11 | 11 | 6 |
| AT1G58080.1 | ATP phosphoribosyl transferase 1 | 10 | 19 | 13 |
| AT3G01500.2 | carbonic anhydrase 1 | 10 | 14 | 11 |
| AT1G49780.1 | plant U-box 26 | 9 | 8 | 11 |
| AT3G06850.1 | 2-oxoacid dehydrogenases acyltransferase family protein | 9 | 9 | 16 |
| ATCG00540.1 | photosynthetic electron transfer A | 9 | 7 | 3 |
| AT1G29920.1 | chlorophyll A/B-binding protein 2 | 8 | 13 | 9 |
| AT5G01600.1 | ferretin 1 | 6 | 3 | 12 |
| AT5G65440.3 | unknown protein | 3 | 4 | 3 |

[a]Accession number of TAIR (http://www.Arabidopsis.org)
[b]Total number of peptide spectrum match (PSM) from 3 independent replicates of WT, AtSAP5-oe-11, and AtSAP5-oe-12

**Table S2.** Proteins identified in WT, AtSAP5-oe-11, and AtSAP5-oe-12 by immunoprecipitation using anti-CAT antibody

| ID[a] | Alias | PSM[b] | Ubiquitination site[c] |
|---|---|---|---|
| **WT** | | | |
| AT4G35090.1 | catalase 2 | 20 | |
| AT1G20620.1 | catalase 3 | 17 | |
| AT5G35790.1 | glucose-6-phosphate dehydrogenase 1 | 13 | |
| ATCG00490.1 | ribulose-bisphosphate carboxylases | 11 | |
| AT3G01670.1 | unknown protein. | 10 | |
| AT2G42600.2 | phosphoenolpyruvate carboxylase 2 | 7 | |
| AT3G09790.1 | ubiquitin 8 | 7 | LIFAGKQLEDGR |
| AT3G01120.1 | Pyridoxal phosphate (PLP)-dependent transferases superfamily protein | 4 | |
| ATCG00480.1 | ATP synthase subunit beta | 4 | |
| AT1G12900.1 | glyceraldehyde 3-phosphate dehydrogenase A subunit 2 | 4 | |
| AT4G27520.1 | early nodulin-like protein 2 | 4 | |
| ATCG00120.1 | ATP synthase subunit alpha | 4 | |
| AT2G02810.1 | UDP-galactose transporter 1 | 4 | TSSKTISK |
| AT2G39730.1 | rubisco activase | 3 | |
| AT3G46520.1 | actin-12 | 3 | |
| AT5G53850.2 | haloacid dehalogenase-like hydrolase family protein | 2 | |
| AT2G28000.1 | chaperonin-60alpha | 2 | |
| AT1G29920.1 | chlorophyll A/B-binding protein 2 | 2 | |
| AT5G54770.1 | thiazole biosynthetic enzyme, chloroplast (ARA6) (THI1) (THI4) | 2 | |
| AT3G54360.1 | zinc ion binding | 2 | |
| AT2G33210.1 | heat shock protein 60-2 | 2 | |
| AT1G13440.1 | glyceraldehyde-3-phosphate dehydrogenase C2 | 2 | |
| AT5G05965.1 | unknown protein | 1 | KGSHTAPEVQGNISQATDERSSHEK |
| AT5G06460.1 | ubiquitin activating enzyme 2 | 1 | |
| AT1G20500.1 | AMP-dependent synthetase and ligase family protein | 1 | QVIDFISKQVAPYKK |
| AT2G40520.4 | Nucleotidyltransferase family protein | 1 | |
| AT5G47110.1 | Chlorophyll A-B binding family protein | 1 | |
| AT3G07610.3 | Transcription factor jumonji (jmjC) domain-containing protein | 1 | |
| AT4G15760.2 | monooxygenase 1 | 1 | |
| AT4G32420.1 | Cyclophilin-like peptidyl-prolyl cis-trans isomerase family protein | 1 | |

| | | | |
|---|---|---|---|
| AT1G60550.1 | enoyl-CoA hydratase/isomerase D | 1 | |
| AT2G34560.2 | P-loop containing nucleoside triphosphate hydrolases superfamily protein | 1 | |
| AT4G26910.1 | Dihydrolipoamide succinyltransferase | 1 | |
| AT4G16630.1 | DEA(D/H)-box RNA helicase family protein | 1 | |
| AT1G67090.1 | ribulose bisphosphate carboxylase small chain 1A | 1 | |
| AT3G01500.2 | carbonic anhydrase 1 | 1 | |
| AT4G23990.1 | cellulose synthase like G3 | 1 | YSPITYGVK |
| AT5G53450.1 | OBP3-responsive gene 1 | 1 | |
| AT1G79150.1 | binding | 1 | |
| AT3G48830.1 | polynucleotide adenylyltransferase family protein / RNA recognition motif (RRM)-containing protein | 1 | |
| **AtSAP5-oe-11** | | | |
| AT1G20620.1 | catalase 3 | 35 | |
| AT4G35090.1 | catalase 2 | 29 | |
| ATCG00490.1 | ribulose-bisphosphate carboxylases | 20 | |
| AT3G01670.1 | unknown protein | 20 | |
| AT5G35790.1 | glucose-6-phosphate dehydrogenase 1 | 16 | |
| AT2G39730.1 | rubisco activase | 13 | |
| AT1G20630.1 | catalase 1 | 13 | |
| ATCG00120.1 | ATP synthase subunit alpha | 12 | |
| ATCG00480.1 | ATP synthase subunit beta | 9 | |
| AT3G26650.1 | glyceraldehyde 3-phosphate dehydrogenase A subunit | 7 | |
| AT1G42970.1 | glyceraldehyde-3-phosphate dehydrogenase B subunit | 7 | |
| AT3G09790.1 | ubiquitin 8 | 7 | LIFAGKQLEDGR |
| AT3G12780.1 | phosphoglycerate kinase 1 | 6 | |
| AT4G38970.1 | fructose-bisphosphate aldolase 2 | 6 | |
| AT1G12900.1 | glyceraldehyde 3-phosphate dehydrogenase A subunit 2 | 6 | |
| AT2G42600.2 | phosphoenolpyruvate carboxylase 2 | 6 | |
| AT2G21330.1 | fructose-bisphosphate aldolase 1 | 5 | |
| AT2G28000.1 | chaperonin-60alpha | 5 | |
| AT5G17920.1 | Cobalamin-independent synthase family protein | 5 | |
| AT5G38410.3 | Ribulose bisphosphate carboxylase (small chain) family protein | 5 | |
| AT3G14420.1 | Aldolase-type TIM barrel family protein | 4 | |

| | | |
|---|---|---|
| AT3G01500.2 | carbonic anhydrase 1 | 4 |
| AT1G55490.1 | chaperonin 60 beta | 4 |
| AT5G53870.1 | early nodulin-like protein 1 | 4 |
| AT1G28290.1 | arabinogalactan protein 31 | 4 |
| AT4G37930.1 | serine transhydroxymethyltransferase 1 | 4 |
| AT4G27520.1 | early nodulin-like protein 2 | 3 |
| AT3G60750.1 | Transketolase | 3 |
| AT1G13440.1 | glyceraldehyde-3-phosphate dehydrogenase C2 | 3 |
| AT1G67090.1 | ribulose bisphosphate carboxylase small chain 1A | 3 |
| AT3G25860.1 | 2-oxoacid dehydrogenases acyltransferase family protein | 2 |
| AT1G20340.1 | Cupredoxin superfamily protein | 2 |
| ATCG00280.1 | photosystem II reaction center protein C | 2 |
| AT1G07940.1 | GTP binding Elongation factor Tu family protein | 2 |
| AT1G29920.1 | chlorophyll A/B-binding protein 2 | 2 |
| AT4G04640.1 | ATPase, F1 complex, gamma subunit protein | 2 |
| AT1G27090.1 | glycine-rich protein | 2 |
| AT5G02500.1 | heat shock cognate protein 70-1 | 2 |
| AT5G53850.2 | haloacid dehalogenase-like hydrolase family protein | 2 |
| AT2G37690.1 | phosphoribosylaminoimidazole carboxylase, putative / AIR carboxylase, putative | 2 |
| AT5G42480.1 | Chaperone DnaJ-domain superfamily protein | 1 |
| AT5G54770.1 | thiazole biosynthetic enzyme, chloroplast (ARA6) (THI1) (THI4) | 1 |
| AT4G24280.1 | chloroplast heat shock protein 70-1 | 1 |
| ATCG00020.1 | photosystem II reaction center protein A | 1 |
| AT5G55070.1 | Dihydrolipoamide succinyltransferase | 1 |
| AT5G35630.3 | glutamine synthetase 2 | 1 |
| AT1G49240.1 | actin 8 | 1 |
| AT3G46520.1 | actin-12 | 1 |
| AT4G32420.1 | Cyclophilin-like peptidyl-prolyl cis-trans isomerase family protein | 1 |
| AT1G18560.1 | BED zinc finger ;hAT family dimerisation domain | 1 |

| | | | |
|---|---|---|---|
| AT2G34560.2 | P-loop containing nucleoside triphosphate hydrolases superfamily protein | 1 | |
| AT4G16630.1 | DEA(D/H)-box RNA helicase family protein | 1 | |
| AT5G09450.1 | Tetratricopeptide repeat (TPR)-like superfamily protein | 1 | |
| AT1G71810.1 | Protein kinase superfamily protein | 1 | |
| AT1G48840.1 | Plant protein of unknown function (DUF639) | 1 | |
| AT1G55040.1 | zinc finger (Ran-binding) family protein | 1 | |
| AT1G11720.2 | starch synthase 3 | 1 | RTTVGSAQK |
| **AtSAP5-oe-12** | | | |
| AT1G20620.1 | catalase 3 | 39 | |
| AT4G35090.1 | catalase 2 | 26 | |
| AT3G01670.1 | unknown protein | 21 | |
| ATCG00490.1 | ribulose-bisphosphate carboxylases | 20 | |
| AT5G35790.1 | glucose-6-phosphate dehydrogenase 1 | 20 | |
| ATCG00480.1 | ATP synthase subunit beta | 10 | |
| AT2G39730.1 | rubisco activase | 9 | |
| ATCG00120.1 | ATP synthase subunit alpha | 9 | |
| AT1G12900.1 | glyceraldehyde 3-phosphate dehydrogenase A subunit 2 | 8 | |
| AT1G28290.1 | arabinogalactan protein 31 | 8 | |
| AT3G26650.1 | glyceraldehyde 3-phosphate dehydrogenase A subunit | 7 | |
| AT2G42600.2 | phosphoenolpyruvate carboxylase 2 | 7 | |
| AT1G20630.1 | catalase 1 | 7 | |
| AT3G01120.1 | Pyridoxal phosphate (PLP)-dependent transferases superfamily protein | 6 | |
| AT1G42970.1 | glyceraldehyde-3-phosphate dehydrogenase B subunit | 6 | |
| AT3G09790.1 | ubiquitin 8 | 5 | LIFAGKQLEDGR |
| AT3G12780.1 | phosphoglycerate kinase 1 | 4 | |
| AT5G53850.2 | haloacid dehalogenase-like hydrolase family protein | 4 | |
| AT2G37690.1 | phosphoribosylaminoimidazole carboxylase, putative / AIR carboxylase, putative | 3 | |
| AT2G28000.1 | chaperonin-60alpha | 3 | |
| AT4G27520.1 | early nodulin-like protein 2 | 3 | |
| AT3G60750.1 | Transketolase | 3 | |

| | | | |
|---|---|---|---|
| AT2G34560.2 | P-loop containing nucleoside triphosphate hydrolases superfamily protein | 3 | |
| AT5G02490.1 | Heat shock protein 70 (Hsp 70) family protein | 3 | |
| ATCG00280.1 | photosystem II reaction center protein C | 2 | |
| AT1G55490.1 | chaperonin 60 beta | 2 | |
| AT1G29920.1 | chlorophyll A/B-binding protein 2 | 2 | |
| AT3G26520.1 | tonoplast intrinsic protein 2 | 2 | |
| AT2G29570.1 | proliferating cell nuclear antigen 2 | 2 | |
| AT3G25860.1 | 2-oxoacid dehydrogenases acyltransferase family protein | 2 | |
| AT3G01500.2 | carbonic anhydrase 1 | 2 | |
| AT3G14415.2 | Aldolase-type TIM barrel family protein | 2 | |
| AT2G33210.1 | heat shock protein 60-2 | 2 | |
| ATCG00270.1 | photosystem II reaction center protein D | 2 | |
| AT1G07940.1 | GTP binding Elongation factor Tu family protein | 2 | |
| AT3G46520.1 | actin-12 | 2 | |
| AT5G47110.1 | Chlorophyll A-B binding family protein | 2 | |
| AT1G13440.1 | glyceraldehyde-3-phosphate dehydrogenase C2 | 2 | |
| AT5G54770.1 | thiazole biosynthetic enzyme, chloroplast (ARA6) (THI1) (THI4) | 2 | |
| AT4G32420.1 | Cyclophilin-like peptidyl-prolyl cis-trans isomerase family protein | 2 | |
| AT5G62300.2 | Ribosomal protein S10p/S20e family protein | 2 | |
| AT1G20500.1 | AMP-dependent synthetase and ligase family protein | 1 | QVIDFISKQVAPYKK |
| AT5G42480.1 | Chaperone DnaJ-domain superfamily protein | 1 | |
| AT4G15760.2 | monooxygenase 1 | 1 | |
| AT5G44020.1 | HAD superfamily, subfamily IIIB acid phosphatase | 1 | |
| AT4G38970.1 | fructose-bisphosphate aldolase 2 | 1 | |
| AT1G27090.1 | glycine-rich protein | 1 | |
| ATCG00020.1 | photosystem II reaction center protein A | 1 | |
| AT2G02810.1 | UDP-galactose transporter 1 | 1 | TSSKTISK |
| AT1G49780.1 | plant U-box 26 | 1 | |

| | | | |
|---|---|---|---|
| AT2G21660.1 | cold, circadian rhythm, and rna binding 2 | 1 | |
| AT1G60550.1 | enoyl-CoA hydratase/isomerase D | 1 | |
| AT5G20980.1 | methionine synthase 3 | 1 | |
| AT1G20060.1 | ATP binding microtubule motor family protein | 1 | |
| AT4G24280.1 | chloroplast heat shock protein 70-1 | 1 | |
| AT5G14660.1 | peptide deformylase 1B | 1 | |
| AT3G55440.1 | triosephosphate isomerase | 1 | |
| AT1G50500.2 | Membrane trafficking VPS53 family protein | 1 | |
| AT1G61400.1 | S-locus lectin protein kinase family protein | 1 | |
| AT3G44840.1 | S-adenosyl-L-methionine-dependent methyltransferases superfamily protein | 1 | |
| AT1G67090.1 | ribulose bisphosphate carboxylase small chain 1A | 1 | |
| AT5G38410.3 | Ribulose bisphosphate carboxylase (small chain) family protein | 1 | |
| AT4G16630.1 | DEA(D/H)-box RNA helicase family protein | 1 | |
| AT4G23990.1 | cellulose synthase like G3 | 1 | |
| AT2G03140.2 | alpha/beta-Hydrolases superfamily protein | 1 | LIYYAQKNKK |
| AT1G48840.1 | Plant protein of unknown function (DUF639) | 1 | |
| AT1G44575.1 | Chlorophyll A-B binding family protein | 1 | |
| AT3G09850.1 | D111/G-patch domain-containing protein | 1 | |

[a]Accession number of TAIR (http://www.Arabidopsis.org)
[b]The number of peptide spectrum match (PSM) in WT, AtSAP5-oe-11, and AtSAP5-oe-12
[c]The ubiquitination sites are shown in red

**Table S3** Relative abundance of CAT1, CAT2, and CAT3 in WT, AtSAP5-oe-11 and AtSAP5-oe-12.

| | Exp. 1 | Exp. 2 | Exp. 3 | Exp. 4 | Mean[a] |
|---|---|---|---|---|---|
| **CAT1** | | | | | |
| WT | 1 | 1 | 1 | 1 | 1±0 |
| AtSAP5-oe-11 | 1.648 | 1.579 | 1.741 | 1.892 | 1.715±0.135** |
| AtSAP5-oe-12 | 1.203 | 1.170 | 1.080 | 1.679 | 1.283±0.269* |
| **CAT2** | | | | | |
| WT | 1 | 1 | 1 | 1 | 1±0 |
| AtSAP5-oe-11 | 1.285 | 1.433 | 1.452 | 2.764 | 1.733±0.69* |
| AtSAP5-oe-12 | 1.291 | 0.974 | 1.293 | 1.440 | 1.249±0.19* |
| **CAT3** | | | | | |
| WT | 1 | 1 | 1 | 1 | 1±0 |
| AtSAP5-oe-11 | 1.419 | 1.215 | 1.116 | 1.800 | 1.387±0.3* |
| AtSAP5-oe-12 | 1.353 | 0.972 | 1.147 | 1.500 | 1.243±0.23* |

[a]Data represent mean ± SD, n = 4 independent experiments (EXP. 1 to EXP. 4) **, $p < 0.05$, *, $p < 0.01$, Student's *t* test compared to WT

**Table S4** Primers used in this study

| Name of primers and probes | Nucleotide sequence | Description |
|---|---|---|
| **Real-time RT-PCR** | | |
| ACT-qF | 5'-GGCAAGTCATCACGATTGG -3' | Detection of *Actin* |
| ACT-qR | 5'-CAGCTTCCATTCCCACAAAC -3' | |
| AtSAP5-qF | 5'-ACCAGCTAAAGTCGTGATTCG-3' | Detection of *AtSAP5* |
| AtSAP5qR | 5'-AGCGGTTTTGTAGTCGTAGC-3' | |
| PR1-qF | 5'- ATCGTCTTTGTAGCTCTTGTAGG-3' | Detection of *PR1* |
| PR1-qR | 5'- AGGCTAAGTTTTCCCCGTAAG -3 | |
| PR5-qF | 5'- GTGTTCATCACAAGCGGCATT-3' | Detection of *PR5* |
| PR5-qR | 5'-GGGAAGCACCTGGAGTCAAT -3' | |
| CAT2-qF | 5'- GATACCGTACCTTTACACCAGAG-3' | Detection of *CAT2* |
| CAT2-qR | 5'- TCCCAAAGACTTATCAGCCTG-3' | |
| **Construction** | | |
| CAT2-ORF-F | 5'-CACCATGGATCCTTACAAGTATCGTC-3' | Construction of pENTR-CAT2-non-stop |
| CAT2-ORF-non-stop-R | 5'-GATGCTTGGTCTCACGTTCAGAC-3' | |
| GFP-ORF-F | 5'-CACCATGGTGAAGACTAATCTTTTTC-3' | Construction of pENTR-GFP-non-stop |
| GFP-ORF-non-stop-R | 5'-CAGCTCGTCCTTCTTGTACAG-3' | |
| NdeI-SUMO-F[a] | 5'-ATTCCATATGTCGGACTCAGAAGTCAATC-3' | Construction of pET-sumo |
| NotI-SUMO-R[a] | 5'ACATGCGGCCGCACCAATCTGTTCTCTGTGAGC-3' | |
| AtSAP5-ORF-F | 5'-CACCATGGCTCAGAGAACGGAGAAG-3' | Construction of pENTR-AtSAP5-non-stop, AtSAP5-cY, or pAtSAP5-CFP |
| AtSAP5-ORF-non-stop-R | 5'-AACTTTGACCATTTTCGCAGC-3' | |
| CAT2-ORF-F | 5′-CACCATGGATCCTTACAAGTATCGTC-3' | Construction of pENTR-CAT2 or nY-CAT2 |
| CAT2-ORF-R | 5′-TTAGATGCTTGGTCTCACGTTC-3' | |
| BamH1-AtSAP5-F[a] | 5′-CCCCGGATCC*A*ATGGCTCAGAGAACGGAGAAG -3' | Construction of AtSAP5-nLuc |
| SalI-AtSAP5-R[a] | 5′-CCCCGTCGACAACTTTGACCATTTTCGCAGC-3' | |
| KpnI-CAT2-F[a] | 5′CCCCGGTACCATGGATCCTTACAAGTATCGTCCAG-3' | Construction of cLuc-CAT2 |
| SalI-CAT2-R[a] | 5′-CCCCGTCGACTTAGATGCTTGGTCTCACGTTC -3' | |
| NPR1-ORF-F | 5′-CACCATGGACACCACCATTGATGGATTC-3' | Construction of pENTR-NPR1-non-stop or pNPR1-GFP |
| NPR1-ORF-non-stop-R | 5′-CCGACGACGATGAGAGAGTTTACG-3' | |

[b]Underlined sequences indicate the sequence recognized by restriction enzyme. The random 4 nucleotides at the 5' end were added as recommended by the manufacturer.

### References and Notes


1. L. Chang *et al.*, Plant A20/AN1 protein serves as the important hub to mediate antiviral immunity. *PLoS Pathogens* **14**, e1007288 (2018).
2. H. Cao, S. A. Bowling, A. S. Gordon, X. Dong, Characterization of an Arabidopsis mutant that is nonresponsive to inducers of systemic acquired resistance. *The Plant Cell* **6**, 1583-1592 (1994).
3. F. H. Wu *et al.*, Tape-Arabidopsis Sandwich-a simpler Arabidopsis protoplast isolation method. *Plant Methods* **5**, 16 (2009).
4. A. Shevchenko, M. Wilm, O. Vorm, M. Mann, Mass spectrometric sequencing of proteins from silver-stained polyacrylamide gels. *Analytical Chemistry* **68**, 850-858 (1996).
5. H. C. Lu *et al.*, A high-throughput virus-induced gene-silencing vector for screening transcription factors in virus-induced plant defense response in orchid. *Molecular Plant-Microbe Interactions* **25**, 738-746 (2012).
6. M. Karimi, D. Inzé, A. Depicker, GATEWAY™ vectors for Agrobacterium-mediated plant transformation. *Trends in Plant Science* **7**, 193-195 (2002).
7. H. Y. Wu *et al.*, AGROBEST: an efficient Agrobacterium-mediated transient expression method for versatile gene function analyses in Arabidopsis seedlings. *Plant Methods* **10**, 19 (2014).
8. H. Aebi, Catalase in vitro. *Methods in Enzymology* **105**, 121-126 (1984).
9. Y. Xiong, A. L. Contento, D. C. Bassham, Disruption of autophagy results in constitutive oxidative stress in Arabidopsis. *Autophagy* **3**, 257-258 (2007).
10. A. L. Contento, Y. Xiong, D. C. Bassham, Visualization of autophagy in Arabidopsis using the fluorescent dye monodansylcadaverine and a GFP-AtATG8e fusion protein. *The Plant Journal* **42**, 598-608 (2005).
11. M. Tian *et al.*, The combined use of photoaffinity labeling and surface plasmon resonance-based technology identifies multiple salicylic acid-binding proteins. *The Plant Journal* **72**, 1027-1038 (2012).
12. Z. Chen, J. W. Ricigliano, D. F. Klessig, Purification and characterization of a soluble salicylic acid-binding protein from tobacco. *Proceedings of the National Academy of Sciences* **90**, 9533-9537 (1993).
